# The molecular mechanism of on-demand sterol biosynthesis at organelle contact sites

**DOI:** 10.1101/2024.05.09.593285

**Authors:** Naama Zung, Nitya Aravindan, Angela Boshnakovska, Rosario Valenti, Noga Preminger, Felix Jonas, Gilad Yaakov, Mathilda M. Willoughby, Bettina Homberg, Jenny Keller, Meital Kupervaser, Nili Dezorella, Tali Dadosh, Sharon G. Wolf, Maxim Itkin, Sergey Malitsky, Alexander Brandis, Naama Barkai, Rubén Fernández-Busnadiego, Amit R. Reddi, Peter Rehling, Doron Rapaport, Maya Schuldiner

## Abstract

Contact-sites are specialized zones of proximity between two organelles, essential for organelle communication and coordination. The formation of contacts between the Endoplasmic Reticulum (ER), and other organelles, relies on a unique membrane environment enriched in sterols. However, how these sterol-rich domains are formed and maintained had not been understood. We found that the yeast membrane protein Yet3, the homolog of human BAP31, is localized to multiple ER contact sites. We show that Yet3 interacts with all the enzymes of the post-squalene ergosterol biosynthesis pathway and recruits them to create sterol-rich domains. Increasing sterol levels at ER contacts causes its depletion from the plasma membrane leading to a compensatory reaction and altered cell metabolism. Our data shows that Yet3 provides on-demand sterols at contacts thus shaping organellar structure and function. A molecular understanding of this protein’s functions gives new insights into the role of BAP31 in development and pathology.

## Introduction

The evolution of organelles was a key event in enabling efficient biophysical isolation of metabolic reactions in eukaryotic cells. However, this necessitated the concomitant formation of ways for organelles to communicate and transfer metabolites to ensure cellular homeostasis. One fundamental mode of communication between organelles is by the creation of areas of proximity between their membranes, termed contact sites^1,2^. Contact sites (in short, contacts) are specialized zones with a unique lipid and protein composition, held by tethering molecules^3^. While it is now clear that all organelles can create contacts^4^, the first ones that were described^5^ and the most well studied ones since, are those that are formed by the largest organelle in the cell, the Endoplasmic Reticulum (ER)^6^.

The proteome of contact sites comprises many proteins that can function in tethering; transfer of lipids, ions, and other small molecules; as well as regulation of contact extent and/or function^7^. Extensive efforts have been undertaken to map the protein repertoire of various contacts^8–11^, uncovering important information about their activity and regulation. In contrast, much less is understood about the unique lipid composition of contacts.

It has been clearly demonstrated that ER contacts are enriched with sterols and sphingolipids that create a specific membrane micro-environment within the continuous membrane of the ER. These subdomains have been dubbed “detergent resistant” or “raft-like”^12–15^ and in ER-mitochondria contacts were shown to contain seven times higher sterols than the surrounding ER membrane^13^. These lipid subdomains are essential for ER compartmentalization and the initiation of cellular actions that are contact regulated, such as apoptosis and autophagy^16–18^. Hence, it is no surprise that these subdomains are important for optimal cellular and organismal function. For example, it was shown that their absence from the ER-mitochondria contact may promote tumor progression^19^ and affect Alzheimer’s disease^20,21^.

Ergosterol and cholesterol, the major sterols in fungi and mammalian cells, respectively, are the products of a multi-step biosynthesis pathway (Figure S1A)^22^. One sterol precursor, Farnesyl-PP, is a metabolite with multiple potential end-products such as heme, dolichol, and ubiquinone^22^. However, once Farnesyl-PP is processed to squalene, it is committed to be converted, by the post-squalene enzymes, into ergosterol or cholesterol (Figure S1A). Following sterol production, which occurs mostly in the ER and on lipid droplets (LD), the majority is immediately transferred to other organelles with the strongest accumulation being at the plasma membrane (PM)^23^. In the PM, sterols serve essential components, necessary for membrane integrity, fluidity, and proper function of multiple membrane proteins^24^. Alternatively, sterols can be stored as Sterol Esters (SEs) in LDs ^25^, which bud from the ER and are crucial for cellular metabolic homeostasis^26^. Hence, sterols are actively removed from the ER through diverse pathways to ensure an overall low level of this molecule in the ER membrane^27^. Despite that, they are still required for ER contact formation and function^12–15^. Thus, a central, unresolved, question in contact site biology is how the sterol-rich lipid subdomains are formed and retained in the sterol-poor environment of the ER.

In this study, we set out to find the ER protein that organizes the sterol-rich subdomains in the ER membrane of the yeast *Saccharomyces cerevisiae*. We found that Yet3 is a pan-ER contact site membrane protein that interacts with the post-squalene ergosterol biosynthesis enzymes, and recruits this synthome, dubbed the ERGosome^28^, to contacts. This leads to a membrane environment enriched in sterols. We demonstrate that there is an inherent balance of sterols between contacts and the PM and thus, Yet3 is also a master regulator of PM lipid subdomains. Consequently, alterations in expression of Yet3 leads to global cellular metabolic changes, which are conserved upon alterations in the levels of its human homologue, BAP31. Our work sheds molecular light on how membrane domains, required for ER contact function, are formed and maintained, and provides clues to the diverse and central roles of BAP31 in human development and health.

## Results

### Yet3 is a pan-ER contact site protein

To identify lipid-organizing proteins, we searched for proteins enriched in more than one contact site using previous datasets for contact site proteomes^8–10^. As expected, we found proteins such as Vps13-family members, VAPs, and LAMs whose function in tethering or lipid transfer was already well defined ^29–33, 47^. Surprisingly, we found one additional protein, of less characterized function, that had a similar distribution, Yet3 (Yeast Endoplasmic reticulum Transmembrane 3). Yet3 is a protein of the ER with three predicted transmembrane domains (TMD), which has two paralogs Yet1 and Yet2^34^. In addition, it is highly conserved to mammals with its human homolog being BAP31 (B-cell receptor Associated Protein of 31kDa) ^34^. BAP31 is also an ER membrane protein, first discovered due to its role in B-cell receptor maturation^35^. It was previously found as an ER-Mitochondria contact (MAM) resident, and it was shown to affect both apoptosis^36–38^ and autophagy^39^; processes known to require specific lipid subdomains containing both sterols and sphingolipids^16–18^. It was also described as a resident of ER-PM contacts^40^ suggesting that also in mammalian cells it is a pan-contact site protein. While BAP31 was suggested to affect many ER pathways such as secretion, quality control, and contact site formation^38,41–44^, its molecular function is still debated and unclear. Hence, we decided to follow up on Yet3 and BAP31 activity and understand their conserved role in ER contacts.

We first set off to verify the results from previous high-throughput work that suggested Yet3 as a resident of several ER contact sites^8,9^. We verified that tagging of Yet3 on its C terminus (C’) preserves its activity (Figure S1B). With this functional fusion protein, we saw that while endogenous expression of Yet3 demonstrates a homogenous distribution in the ER, it becomes concentrated in specific ER subdomains (reminiscent of contacts) when overexpressed. This was distinct from its paralog, and heterodimer partner, Yet1 (Figure S1C and Figure S1D).

To map the extent of contact sites to which Yet3 resides, we assayed its co-localization with split-Venus reporters for ER contacts with various organelles^45,46^. To create the reporters, we tagged an abundant ER membrane protein, Snd3, with the N’ portion of a split-Venus (Snd3-VN) and attached the other half of the Venus protein (VC) to abundant membrane proteins on opposing organelles: Pex25-VC for peroxisomes; Faa4-VC for LDs; Ina1-VC for PM; Tom20-VC for mitochondrion and Zrc1-VC for the vacuole. We have previously demonstrated that only in areas where contacts are formed (30-80nm), the two parts of the split-Venus protein are close enough to interact in trans, thus emitting a signal that allows visualization of contacts by fluorescence microscopy ^46^. We found that Yet3-mScarlet foci partially co-localized with every contact site reporter that we assayed, suggesting that Yet3 is found in multiple ER contacts (Figure 1A).

**Figure 1.**
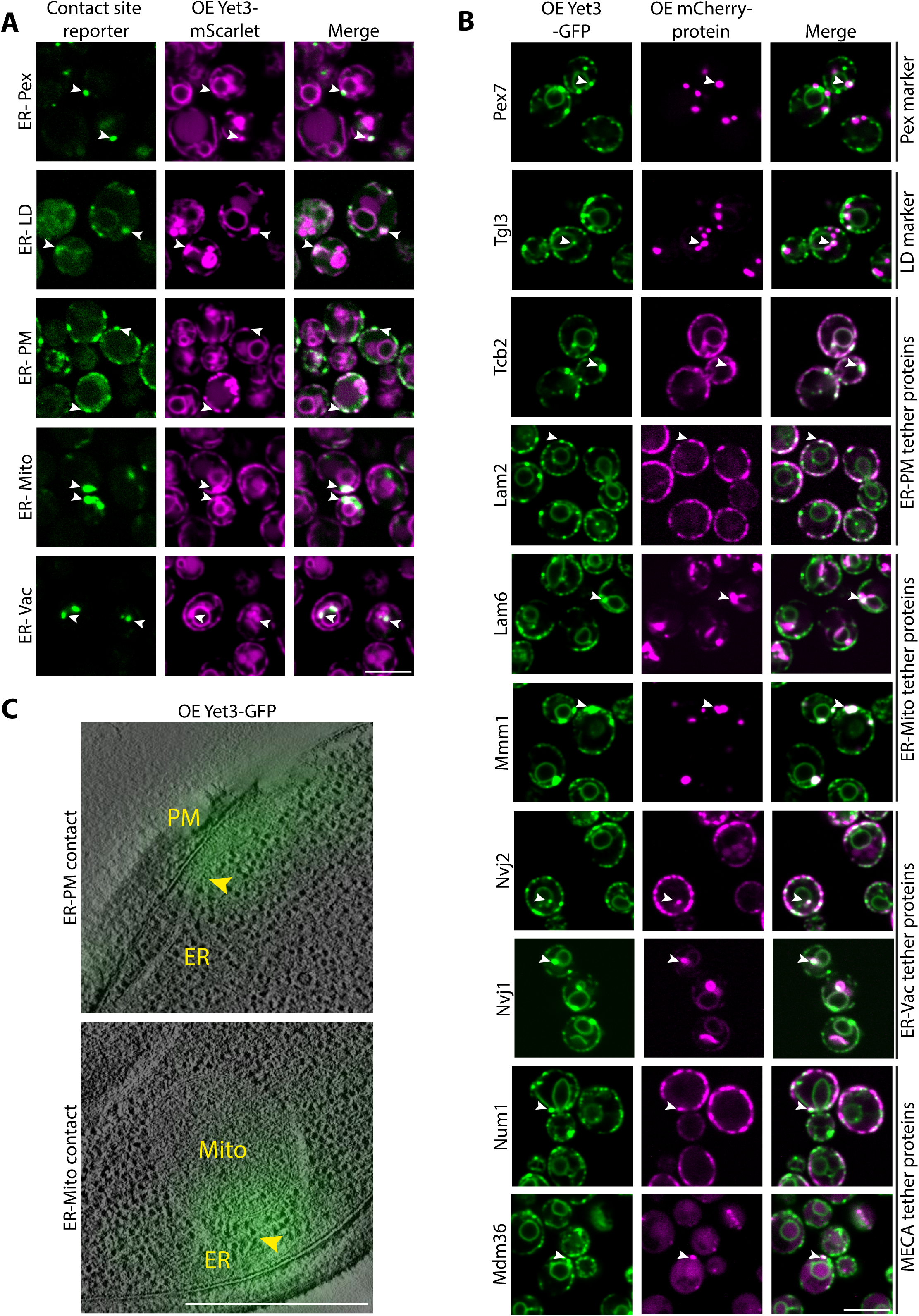
– Yet3 is a pan-ER contact site protein. A. Overexpressed (OE) Yet3-mScarlet concentrates at many ER contact sites. To visualize the localization of Yet3 at contacts using fluorescence microscopy, various split-Venus reporters were used: ER-Peroxisomes (Pex) reporter (Snd3-VN/Pex25-VC); ER-Lipid droplets (LD) reporter (Snd3-VN/Faa4-VC); ER-Plasma membrane (PM) reporter (Snd3-VN/Ina1-VC); ER-Mitochondria (Mito) reporter (Snd3-VN/Tom20-VC) and ER-Vacuole (Vac) reporter (Snd3-VN/Zrc1-VC). White arrows show areas of co-localization between Yet3 puncta and the indicated reporter signals. Strains were imaged with a 100x oil lens. Scale bar, 5µm. B. Overexpressed (OE) Yet3-GFP localizes to contact sites in the absence of a synthetic reporter. Marker proteins (Pex7 and Tgl3 for Pex or LDs, respectively), and tether proteins (Tcb2 and Lam2 in the ER-PM contact, Lam6 and Mmm1 in the ER-Mito contact, Nvj1 and Nvj2 in the ER-Vac contact, Num1 and Mdm36 in the ER-mitochondria-plasma membrane (MECA) contact), were expressed under a constitutive promoter and tagged with mCherry on their N’. White arrows mark areas of co-localization between the Yet3 puncta and the mCherry tagged proteins. Strains were imaged with a 100x oil lens. Scale bar, 5µm. C. Correlated Light and Electron Microscopy (CLEM) images showing that Yet3 localizes to both ER-mitochondria (Mito) and ER-plasma membrane (PM) contacts. The yellow arrows point to the contact site between the ER tubule and the indicated organelle, where the Yet3-GFP signal is bright. Scale bar, 500nm.

While the synthetic reporters do not cause formation of aberrant contact sites^46^, they do affect contact dynamics. Hence, to verify that Yet3 is localized to endogenous contacts without relying on reporters, we assayed contact site co-localization using known tethers for most organelles, or organelle markers for peroxisomes and LDs (Pex7-mCherry and Tgl3-mCherry, respectively) (Figure 1B). We used these organelle markers since imaging contacts in the absence of a reporter is more challenging for peroxisomes and LDs, the size of which (∼100nm in diameter) is below the resolution of conventional light microscopes (∼250nm). At this resolution it is not possible to say if the protein indeed resides in the contact or merely co-localizes with the entire organelle. For all other ER contacts, we visualized known tethers such as Tcb2 and Lam2^47,48^ for ER-PM; Lam6 and Mmm1^30,33,49^ for ER-Mitochondria, Nvj1 and Nvj2^50,51^ for ER-Vacuole, and Num1 and Mdm36 for the three-way contact site between the Mitochondria-ER-Cortex (PM), also known as the MECA^52^. We found that they all indeed partially co-localized with Yet3 puncta (Figure 1B).

To further verify and visualize the presence of Yet3 in contacts at a higher resolution, we performed correlative light and electron microscopy (CLEM). CLEM enables easy detection of ER-mitochondria and ER-PM contacts, and indeed we found overexpressed Yet3-GFP concentrates abundantly at these two contacts (Figure 1C).

Put together, our results demonstrate that Yet3 is a pan-ER contact membrane protein.

### Yet3 levels affect organelles in an Opi1-independent manner

Contact residents can often influence the opposing organelles^9,49^. Therefore, we set out to find how Yet3 overexpression affects the organelles with whom the ER makes contact. To visualize the various organelles, we C’ tagged with mCherry either Pex3 (peroxisomes), Tgl3 (LDs) or Tom20 (mitochondria). For vacuole membrane visualization, we used the FM_4-64_ dye. We found that all organelles tested were affected by the high levels of Yet3: peroxisome numbers increased significantly, LD numbers were significantly reduced, vacuoles were enlarged, and mitochondrial shape was altered (Figure 2A and Figure S2A).

**Figure 2.**
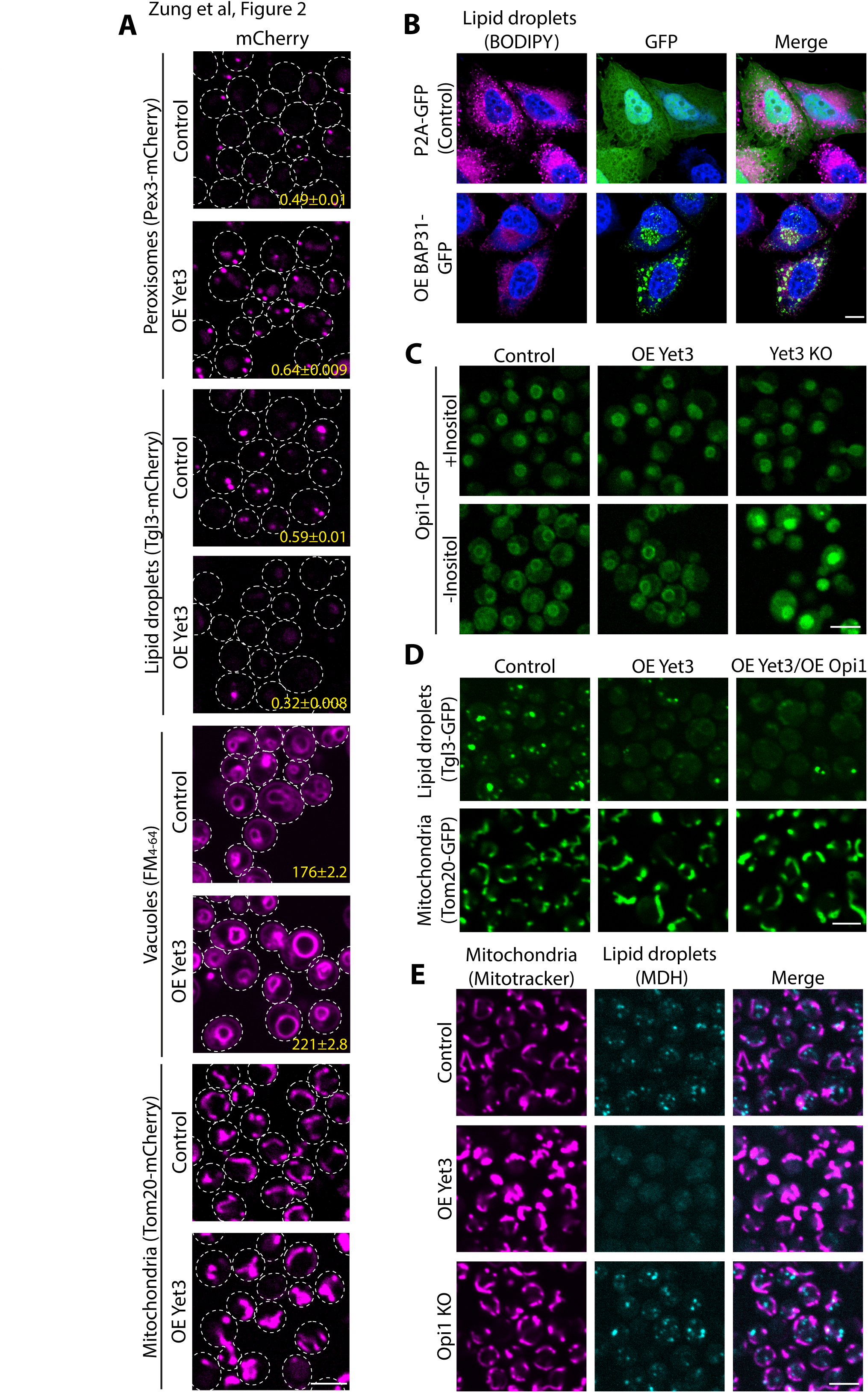
– Yet3 levels affect organelles in an Opi1-independent manner. A. Microscopy images highlighting how multiple organelles are affected by overexpression (OE) of Yet3. In strains that overexpress Yet3, peroxisome number (marked using Pex3-mCherry) increased, while LD number (marked with Tgl3-mCherry) decreased. The vacuoles (dyed with FM_4-64_) and mitochondria (shown by Tom20-mCherry) appear enlarged. The mean number of peroxisomes and LDs per cell was quantified and is presented in yellow at the bottom of each image, with standard error of mean. The mean of the vacuole area per cell was quantified and is presented in yellow at the bottom of the image, with standard error of mean. The differences were statistically significant using a two-tailed t-test, ****p ≤ 0.0001. In the peroxisome analysis, N= 5188, 7090 for control and OE Yet3 respectively. In the LD analysis, N=5648, 6378 for control and OE Yet3 respectively. In the vacuole analysis, N= 6734, 6378 for control and OE Yet3 respectively. Cells were imaged with a 60x oil lens. Scale bar, 5µm B. HeLa S3 cells display a decreased number of LDs following overexpression (OE) of the human homolog of Yet3, BAP31. HeLa S3 cells were transfected with either P2A-GFP plasmid as a control or BAP31-GFP plasmid for overexpressing BAP31. LDs were visualized using BODIPY red, and Hoechst dye was used for nuclear staining in blue. Shown are representative images from three replicates. Cells were imaged using a 63x glycerol lens. Scale bar, 10µm. C. Increased expression of Yet3 does not cause mis-localization of Opi1. On the background of Yet3 overexpression (OE) or knockout (KO), Opi1-GFP consistently enters the nucleus when inositol is present. Depletion of inositol from the media showed an Opi1-GFP signal on the nuclear membrane in both control and Yet3 overexpression. However knockout of Yet3 led to Opi1 accumulating inside the nucleus as expected. All strains were imaged with a 60x oil lens. Scale bar, 5µm. D. Increased levels of Opi1 in inositol-containing media did not rescue the phenotypes of overexpressed (OE) Yet3. Mitochondria were visualized by Tom20-GFP and LDs using Tgl3-GFP. Images were taken using a 60x oil lens. Scale bar, 5µm. E. Knockout (KO) of *OPI1* does not mimic the Yet3 overexpression (OE) phenotypes. Mitochondria were imaged using Mitotracker Orange and LDs by using the blue MDH dye. Images were taken using a 60x oil lens. Scale bar, 5µm.

To test if this effect is conserved to BAP31, we overexpressed BAP31-GFP in HeLa S3 cells. While BAP31 was shown to be homogenously distributed in the ER^53,54^, its overexpression led to its concentration at specific subdomains in the ER, similar to the Yet3 foci observed in yeast (Figure 2B). Additionally, the number of LDs as visualized with BODIPY red, was dramatically reduced (Figure 2B), suggesting that BAP31’s cellular effect is conserved from yeast to humans.

How can Yet3 expression levels cause such global cellular effects? Previous work showed that Yet3 and its paralog Yet1, create a heterodimeric complex that regulates the inositol biosynthesis pathway by binding the master regulator of inositol biosynthesis, Opi1^55^. In summary, when inositol is present, Opi1 hinders inositol synthesis by entering the nucleus and inhibiting Ino2/Ino4 transcriptional activation, causing reduced expression of both inositol and phospholipid biosynthesis enzymes such as Ino1 and Cho2 (Figure S2B). In contrast, inositol depletion causes Opi1 to bind the Yet1-Yet3 heterodimer together with the yeast VAP, Scs2, on the nuclear membrane. This inhibits Opi1 entrance into the nucleus and enables the activation of all Ino2/Ino4 transcriptional targets (Figure S2B). *Yet3* or *yet1* deletion, therefore, disturbs the tethering of Opi1 to the nuclear-ER in case of inositol depletion from the media, and could cause dramatic cellular rewiring. Yet3 overexpression, conversely, could cause Opi1 to be consistently sequestered to the nuclear envelope, even when inositol is present, thereby altering its capacity to downregulate inositol production. Indeed, all our experiments thus far were performed in inositol-rich media, a condition under which it could be deleterious to prevent Opi1 from entering the nucleus.

To test if excessive or dysregulated inositol biosynthesis is the direct reason for the organelle number and morphology changes observed, we imaged Opi1-GFP while manipulating the expression levels of Yet3. In media depleted of inositol, Opi1 was found on the nuclear membrane in either control or overexpression of Yet3, as expected. Moreover, Opi1 remained inside the nucleus in a Δ*yet3* background, as was previously reported (Figure 2C)^55^. However, increased levels of Yet3 did not affect nuclear Opi1 localization in media containing inositol (Figure 2C). This may be because Yet1 levels remain unchanged and hence the heterodimeric complex levels do not increase. Regardless of the mechanistic explanation, this suggested that mis-localization of Opi1 is not the underlying reason for the phenotypes that we observed. This also prompted us to rely more on Yet3 overexpression from here on forward.

To verify that the phenotypes are not an indirect effect of Opi1 sequestration, we examined whether overexpression of Opi1 (which should lead to free Opi1 capable of entering the nucleus^56^) can rescue Yet3 overexpression phenotypes. Using LD number and mitochondrial shape as our phenotypic readout, we found that increased expression of Opi1 could not rescue these phenotypes (Figure 2D). Furthermore, when we imaged an *opi1* mutant, which would mimic a state of increased Opi1 tethering to the nuclear membrane –leading to elevated concentration of inositol in the cell^56^ (Figure S2B), we found that it did not imitate the effect of overexpressing Yet3 (Figure 2E).

Put together, our results all demonstrate that Yet3 has an Opi1-independent role whose expression levels dramatically alter organelles architecture.

### Yet3 interacts with the post-squalene ergosterol biosynthesis machinery affecting sterol distribution in the cell

If Yet3 is not working through its heterodimeric role with Yet1 to sequester Opi1, what is it doing at contact sites and how does it cause such a broad cellular effect when overexpressed? To investigate the additional role of Yet3, we identified its interactors using immunoprecipitation followed by Mass Spectrometry (IP-MS). Interestingly, we found that overexpressed Yet3-GFP showed an enrichment of all the post-squalene ergosterol biosynthesis proteins (Figure 3A, Figure S1A) (Table S1)^22^. This suggested that Yet3 somehow affects the committed step in sterol biosynthesis at contact sites.

**Figure 3.**
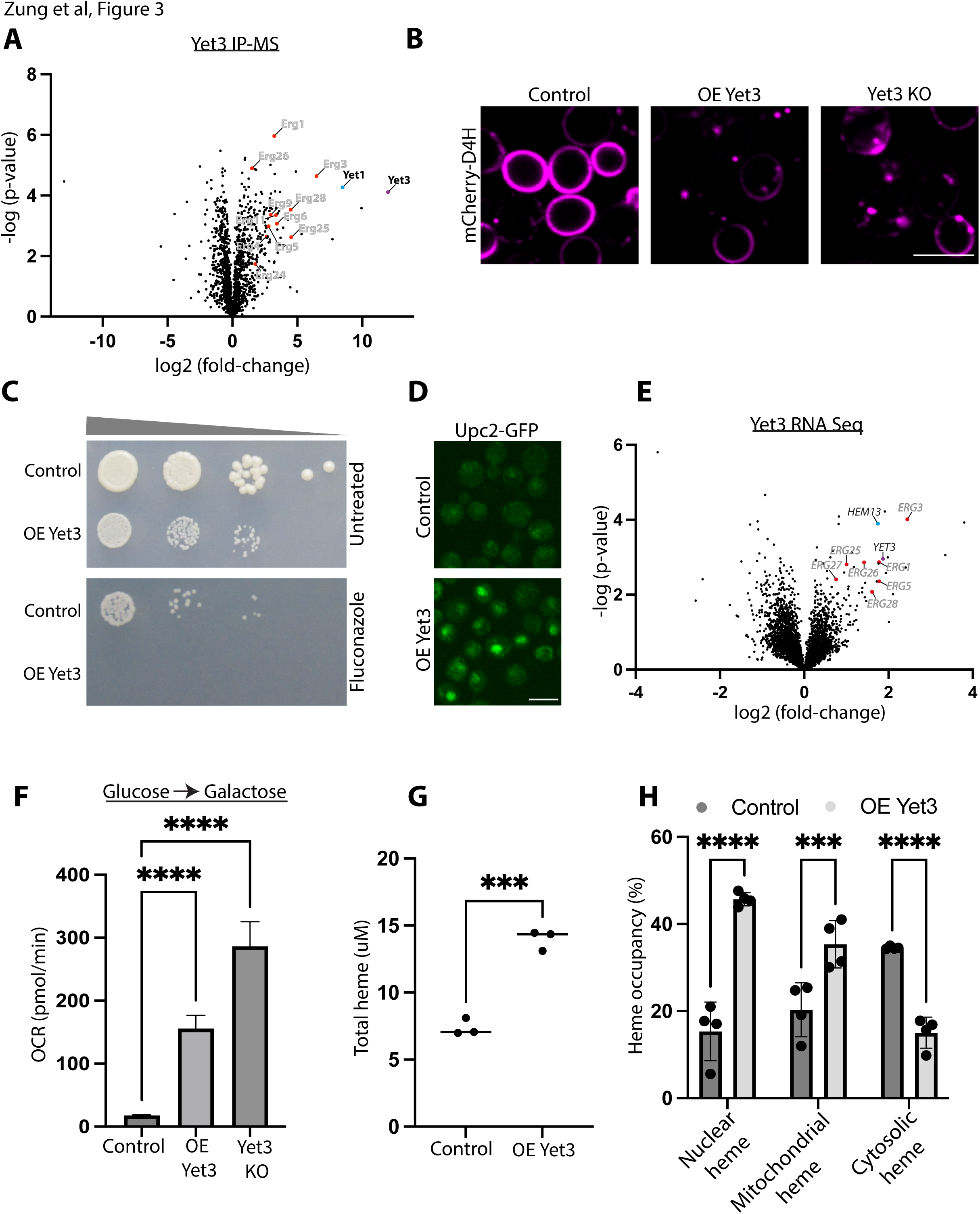
– Yet3 interacts with the post-squalene ergosterol biosynthesis machinery, affecting sterol distribution in the cell. A. Ergosterol biosynthesis proteins are enriched as interactors of Yet3. A volcano plot showing – log(p-value) vs. log2(fold-change) of changes observed following Immunoprecipitation-Mass Spectrometry (IP-MS) to identify peptides of interacting proteins when compared to an overexpressed Tom20-GFP control. Highlighted are enriched interactors of overexpressed Yet3-GFP. Ergosterol biosynthesis proteins, shown in red, are both enriched and are statistically significant (*p ≤ 0.05). Yet3 itself is represented by a purple dot, while Yet1 is represented in a blue dot. Shown are average enrichment values from biological triplicates. B. Yet3 expression levels affect the distribution of plasma membrane sterols. mCherry-D4H, a reporter for free ergosterols, was expressed on the background of control, overexpressed (OE) Yet3 and Yet3 knockout (KO) strains, demonstrating altered sterol distribution in the cell. All samples were imaged with a 100x oil lens. Scale bar, 5µm. C. Overexpression (OE) of Yet3 sensitized cells to the ergosterol-biosynthesis inhibiting drug, fluconazole. Drop dilution assay of control and overexpressed Yet3 strains grown on control (Untreated) or Fluconazole (20ug/ml) containing media. Images were acquired after three days at 30°C. A representative image from three replicates is shown. D. The transcriptional activator of ergosterol biosynthesis, Upc2, enters the nucleus upon overexpression (OE) of Yet3. Upc2, known to translocate from the cytosol to the nucleus upon sensing a reduction in PM ergosterols, was tagged with GFP on its C’ to visualize its cellular localization by fluorescence microscopy using a 60x oil lens. Scale bar, 5µm. E. mRNAs of ergosterol biosynthesis enzymes are upregulated in strains overexpressing Yet3 compared to control cells. The volcano plot shows –log(p-value) against log2(fold change) of changes recorded using RNA-Seq. Highlighted in red are the post-squalene pathway transcripts, *HEM13* is marked in blue and the *YET3* mRNA is in purple. Next Generation Sequencing (NGS) was performed in three replicas per strain. F. Oxygen consumption rate (OCR) increases in overexpression (OE) of Yet3 and in the Yet3 knockout (KO). All samples were grown on glucose overnight and transferred to galactose for 24 hours before measuring their basal respiration. Significance of the changes for three independent replicates was tested using two-way ANOVA. ****p ≤ 0.0001. G. Total heme concentration in overexpressed (OE) Yet3 is higher than in control. A porphyrin fluorescence assay was used to measure heme concentration. Significance of the changes for three independent replicates was tested using an unpaired t-test. ***p ≤ 0.001. H. In contrast to mitochondria and the nucleus, cytosolic free heme was reduced in overexpression (OE) of Yet3. Unbound heme concentration was measured using the Heme Sensor1 (HS1). HS1 levels of heme were measured by heme dependent fluorescence emission compared to an independent fluorophore by percentage. The HS1 was targeted to the mitochondrial matrix by fusing it to the N’ of Cox4, or to the nucleus by fusing it to the C’ of SV40. Significance of the changes from three independent replicates was tested using two-way ANOVA significance test. ***p ≤ 0.001. ****p ≤ 0.0001.

To further explore the connection between Yet3 overexpression and ergosterol, we followed free ergosterol distribution using the free-sterol biosensor, D4H^57^, tagged with mCherry. In the control strain, we found the D4H signal mainly on the PM as previously reported^57^. Interestingly, both Yet3 overexpression and deletion led to reduced PM staining and re-localization of D4H to internal foci (see discussion) (Figure 3B).

Since there was an effect on sterol distribution at the PM, we assayed how Yet3 levels affect cell growth in the presence of the antifungal drug fluconazole. Fluconazole inhibits Erg11, an essential enzyme in the post-squalene ergosterol biosynthesis pathway, which leads to reduction of ergosterol levels in the cell (Figure S1A). Since the majority of ergosterols are transferred to the PM^23^, it would also be the membrane most affected by fluconazole addition^58^. Indeed, the strain overexpressing Yet3 was susceptible to fluconazole (Figure 3C). It was also shown that reduced sterols increase the levels of long-chain sphingolipids as a compensatory mechanism^59^. Indeed, lipidomic analysis demonstrated reduced sterol levels in strains overexpressing Yet3 (Figure S3A), and elevated levels of long-chain sphingolipids (Figure S3B) (Table S2).

Reduced sterols on the PM should cause mis-localization of PM proteins that depend on sterol-rich subdomains for their localization. Examples of such proteins are Gap1 and Agp1, two broad-range amino acid permeases^60–62^. Indeed, Gap1-GFP and Agp1-GFP are both mis-localized on the background of *yet3* deletion and Yet3 overexpression (Figure S3C). Polar metabolite profiling analysis further supports this change in PM permeases since the levels of several amino acids are reduced in the overexpressed Yet3 strain compared to the control (Figure S3D) (Table S3). In addition, the amino acid responsive transcription factor, Gcn4, translocate to the nucleus, indicative of reduced overall amino acid levels (Figure S3E)^63^.

Another known outcome of sterol reduction on the PM, is the activation of the ergosterol sensor and transcription factor Upc2^64^. Loss of PM sterols causes Upc2 to reveal a nuclear localization sequence, resulting in its nuclear accumulation and upregulation of transcripts encoding for ergosterol biosynthesis enzymes^64^. Indeed, Upc2-GFP shifted its localization from cytosolic to nuclear when Yet3 was overexpressed (Figure 3D). In addition, transcriptomic analysis by RNA-Seq verified that cells overexpressing Yet3 have increased levels of Upc2 and its targets, the post-squalene enzyme transcripts, compared to control cells (Figure 3E) (Table S4).

If the increased mRNAs for all the post-squalene biosynthetic enzymes indeed cause elevated ERG protein production and function, then also oxygen consumption rates should increase since some enzymes in the pathway require oxygen, such as Erg11, Erg3 and Erg5 (Figure S1A). To see if oxygen consumption levels are affected by Yet3, we measured oxygen consumption rate (OCR) via Seahorse for different Yet3 expression levels (Table S5). Strains were grown in glucose, and then transferred to galactose containing medium for the analysis. Interestingly, both the Yet3 overexpressed strain and the deletion strain of *yet3* showed an increase in OCR compared to the control (Figure 3F). OCRs were most likely not elevated due to increased cellular respiration since translation of mitochondrial transcripts encoding the electron transport chain subunits were decreased in overexpression of Yet3 (Figure S3F); and mitochondrial respiration during growth in the non-fermentable carbon source, ethanol, was unchanged in overexpressed Yet3, and only slightly elevated in Δ*yet3* (Figure S3G). Altogether our data suggest that mitochondrial respiration is not upregulated, and that instead the increased OCR may result from oxygen shunted for sterol production.

Sterol production also requires large amounts of the carbon precursor, acetyl-CoA. In support for central carbon metabolism being shunted to provide this precursor in strains overexpressing Yet3, RNAseq results show that central carbon metabolism is downregulated at the transcriptional level (Table S4). Moreover, polar metabolite profiling analysis uncovered a reduction in the intermediate metabolites of glycolysis (Figure S3H) and tricarboxylic acid (TCA) cycle (Figure S3I) (Table S3), with a concomitant increase in pyruvic acid, the exit metabolite of glycolysis, and citrate, the entry metabolite of the TCA cycle (Figure S3H and Figure S3I). Indeed, pyruvic acid is important for acetyl Co-A production^22^. Moreover, two of the four amino acids whose amount increased upon overexpression of Yet3, alanine and arginine (Figure S3D), are important for CoA synthesis, which is the precursor for acetyl CoA^65,66^.

We next tested heme levels, since this is an essential co-factor of several enzymes of the post-squalene pathway. Our RNA-Seq data showed increased transcript levels of *HEM13,* which encodes the heme biosynthetic enzyme coproporphyrinogen oxidase, in the Yet3 overexpression strain (Figure 3E). To measure total heme concentrations more directly, we used a porphyrin fluorescence assay. This assay revealed elevated overall heme levels in cells overexpressing Yet3 as would be expected from the higher levels of Hem13 (Figure 3G). Further, we used the genetically encoded ratiometric fluorescent heme sensor HS1-M7A to assess free heme levels^67^. The fractional heme occupancy of HS1-M7A is proportional to the heme available to the sensor. Free heme in the cytosol was diminished, despite the nuclear and mitochondrial fractions being elevated (Figure 3H). These results are consistent with an increased utilization of heme by the ergosterol biosynthesis enzymes that face the cytosol.

Altogether, our data demonstrate that Yet3 interacts with the post-squalene enzymes and affects sterol distribution in the cell. Despite the increase in heme levels, oxygen consumption, and *ERG* gene transcription – PM sterols remain low. This raises the question of whether the produced ergosterol is somehow sequestered by Yet3 in ER contacts.

### Yet3 recruits the ERGosome to provide on-demand sterol biosynthesis at ER contact sites

Since Yet3 interacts with all the post-squalene pathway enzymes and affects sterol distribution, we hypothesized that it plays a role in sequestering the ergosterol enzymes at contact sites. To this end, we tagged 13 out of the 15 post-squalene ergosterol biosynthesis enzymes with mCherry on their N’ (tagging that allowed their clear visualization at their native locations^68^) and imaged the effect of Yet3 expression on their localization. Strikingly, we found that upon overexpression of Yet3, the localization of all 13 tagged enzymes was shifted from their homogenous distribution on either the ER or in LDs, to co-localize with Yet3-GFP puncta (Figure 4A and Figure S4A). This was in sharp contrast to the pre-squalene enzymes Hmg1/2 whose localization was not altered upon overexpression of Yet3 (Figure 4A and Figure S4A). Our data suggest that Yet3 sequesters, directly or indirectly, all the post-squalene biosynthesis enzymes to create a “synthome”, previously dubbed the ERGosome^28^, for on-demand ergosterol biosynthesis at contact sites. The ERGosome was also observed as a potential high molecular weight complex harboring Yet3 by blue native gel electrophoresis (Figure S4B). This is further supported by previous complexome analysis of mitochondrial proteins where Yet3 associated with a complex of approximately 750kDa also shared by Erg4 and Erg26^69^.

**Figure 4.**
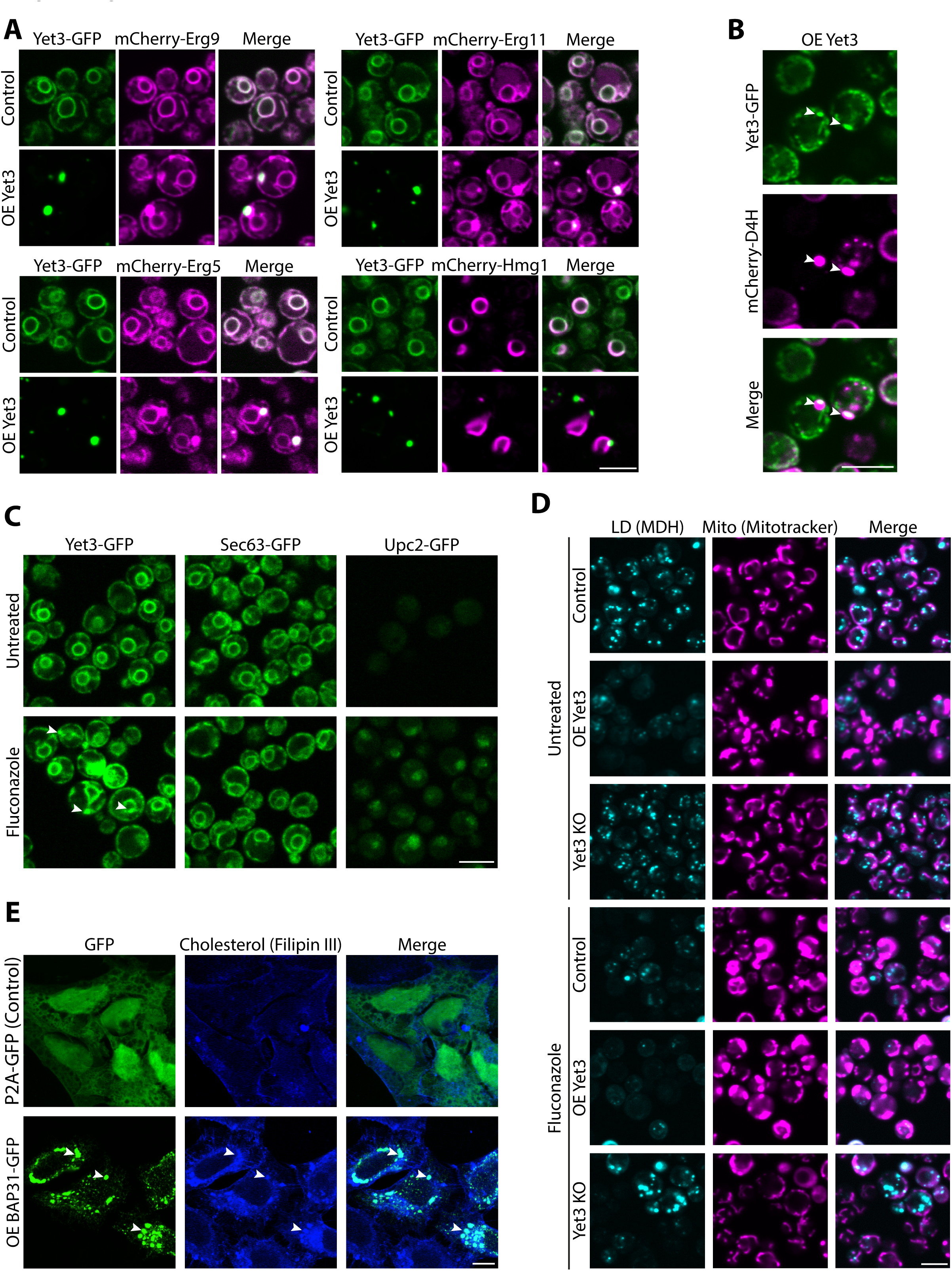
– Yet3 recruits the ERGosome to provide on-demand sterols at ER contact sites. A. Yet3 overexpression (OE) alters the localization of post-squalene biosynthesis pathway proteins to specific subdomains on the ER membrane. Constitutively expressed post-squalene proteins tagged with mCherry on their N’ were homogeneously distributed in strains expressing endogenous Yet3-GFP. Upon overexpression of Yet3, they concentrated in ER subdomains that co-localize with Yet3 puncta. The pre-squalene enzyme, mCherry-Hmg1, did not alter its distribution nor concentrate together with overexpressed Yet3 puncta, which demonstrates that this is not an unspecific restructuring of the ER as a whole. Images were taken with a 100x oil lens. Scale bar, 5µm. B. Yet3-GFP containing foci co-localized with sterol-rich areas (measured using the free ergosterol reporter mCherry-D4H), showcasing that internal ergosterol accumulates at contact sites when Yet3 is overexpressed. Strains were imaged with a 100x oil lens. Scale bar, 5µm. C. During conditions of decreased ergosterol abundance in the cell (created by treating the cells with the ergosterol biosynthesis inhibitor Fluconazole), Yet3, expressed under its own promoter, accumulates at ER subdomains. Endogenously expressed Yet3-GFP is homogenously distributed around the ER. Applying Fluconazole (20ug/ml), caused sterol depletion from cells as can be seen by the translocation into the nucleus of Upc2-GFP, a transcription factor that senses ergosterol reduction in the PM and enters the nucleus. Under these conditions, Yet3-GFP also accumulates in subdomains of the ER, while another ER protein, Sec63-GFP, does not change. Strains were imaged in 60x lens. Scale bar, 5µm. D. Yet3 depletion rescues Fluconazole induced cellular alterations. Yeast strains with endogenous (Control), overexpressing (OE) or knocked out (KO) for Yet3 were imaged in regular media or treated with Fluconazole (20ug/ml) to inhibit ergosterol biosynthesis. Fluconazole addition reduced LD number and altered the shape of mitochondria (Mito) in a way that pheno-mimicked the Yet3 overexpression alone. Combined overexpression of Yet3 and growth in fluconazole led to enhanced phenotypes. Importantly, knockout of Yet3 rescued both LD abundance and mitochondrial morphology in cells treated with Fluconazole compared to untreated control cells. Mitochondria were stained with Mitotracker Orange, and LDs were stained using MDH. Strains were imaged in PBS. Scale bar, 5µm. E. BAP31 overexpression alters cholesterol distribution. Confocal images of HeLa S3 cells transfected with an overexpression (OE) BAP31-GFP plasmid, demonstrate changes in cholesterol (visualized using Filipin III dye) distribution in the cell. P2A-GFP plasmid was transfected to HeLa S3 as control. White arrows in cells overexpressing BAP31 indicate co-localization between cholesterol concentrations and BAP31 puncta on the ER. Cells were imaged using a 63x glycerol lens. Scale bar, 10µm.

If indeed Yet3 recruits the ERGosome to contact sites, we would also expect to see ergosterol accumulating at Yet3 foci. In support of this, using the mCherry-D4H reporter, we observed that free ergosterols were mainly co-localized with Yet3 containing foci when Yet3 was overexpressed (Figure 4B).

Previous reports showed that Yet3 heterodimerizes with its paralog Yet1, for inositol biosynthesis regulation^55,70^. Although we already eliminated inositol biosynthesis regulation as the pathway by which Yet3 affects contact sites (Figure 2), it could still be that its heterodimerization with Yet1 is important for its role in ERGosome scaffolding. However, our data supports a Yet1-independent role for Yet3 for three reasons. First, we observed Yet3 is much more abundant than its paralogs (Yet1 or Yet2), suggesting heterodimer-independent activity (Figure S1D and S4C). Second, blue native gel analysis indicated that deleting *yet1* does not dramatically affect the high molecular weight complex assembly (Figure S4B). Third, deletion of *yet1* increased the number and size of overexpressed Yet3 subdomain puncta and caused elevated ergosterol levels in the contacts, as shown by D4H, while overexpressing *YET1* reduced Yet3 puncta and decreased the amount of ergosterols in the contacts (Figure S4D). Hence, Yet1 does not support the role of Yet3 in ERGosome scaffolding, but rather it may sequester Yet3 for its role in Opi1 localization, thus regulating its capacity to translocate to contact sites and induce local sterol production.

So far, our data demonstrate that overexpression of Yet3 causes it to accumulate at contacts, initiating scaffolding of the ERGosome and increasing ergosterol levels at contacts at the expense of their trafficking to the PM. While this suggests that Yet3 is sufficient to induce local sterol production, it does not clearly demonstrate that it is necessary for this process when expressed under its own endogenous promoter. We therefore examined whether endogenous Yet3-GFP is also recruited to contacts. We hypothesized that Yet3 will be recruited to contacts during ergosterol deficiency, in other words, when ergosterol is in demand. Indeed, when grown in the presence of fluconazole, resulting in an ergosterol-depleted environment (shown by accumulation of Upc2 in the nucleus), we observed Yet3 (Figure 4C) and the ergosterol biosynthesis proteins (Figure S4E) concentrated at subdomains. This was also seen in iron-depleted medium that should cause reduction in free ergosterol due to reduced heme production (Figure S4F).

To support our assumption that native Yet3 is necessary for “on demand” sterol biosynthesis at the contacts, we relied on the fact that addition of fluconazole pheno-mimicked the organellar changes observed when Yet3 was overexpressed: fewer LDs and bulkier mitochondria (supported by a report that showed mitochondrial morphology changes when the ergosterol pathway is blocked^71^) (Figure 4D). The fact that the combination of growth in fluconazole alongside overexpression of Yet3 exacerbated the phenotypes supports the idea that each of these changes cause an independent reduction in total ergosterol. Importantly, deletion of *yet3* rescued the phenotypes of fluconazole treatment (Figure 4D). This evidence strongly supports that under endogenous conditions, the presence of Yet3 causes a certain fraction of ergosterol to be sequestered at contacts. Freeing this fraction by deletion of *yet3* releases the ergosterols from contact sites to other metabolic uses in the cell, thus rescuing the overall phenotypic consequences of reduced sterol levels.

If on-demand sterol biosynthesis at contact sites is a central function of Yet3, then we would expect it to be conserved to its human homolog BAP31. Indeed, cholesterol was concentrated more in the ER than on the PM in BAP31 overexpression compared to the control (Figure 4E).

Taken together, our results uncover a novel mechanism that orchestrates the formation and maintenance of lipid subdomains at contact sites. Furthermore, the role of Yet3/BAP31 may elucidate the extensive involvement of BAP31 in numerous pathways, many of which are dependent on sterol enriched subdomains. This, in turn, provides a new perspective on its potential impact on a multitude of developmental processes and in disease states.

## Discussion

In recent years it has become apparent that the ER membrane has subdomains with unique lipid compositions. One type of such subdomain consists of sterol-rich areas that are required for the formation of contact sites. Additional functions of ER subdomains have been demonstrated such as during fusion of tubules required for ER shaping, translocation of unique substrates, and trafficking of special cargo at ER exit sites^72–75^. While the proteome of ER contact sites was extensively studied over the last decade, how the distinctive lipid composition of these subdomains is formed and maintained, had remained elusive.

In this study, we uncovered the molecular mechanism that enables the formation of unique lipid domains enriched with sterols at ER contact sites. Ergosterol is the final product of a multi-step enzymatic biosynthesis pathway (Figure S1A). Once the dedicated intermediate, squalene, is formed – the pathway is made efficient by organization of the post-squalene enzymes as a synthome, dubbed the ERGosome^28^. Here, we show that the ERGosome is recruited to ER contact sites by Yet3, an ER membrane spanning protein. Moreover, this recruitment underlies the formation of the sterol-rich environment of contacts (Figure 5). We observed that while Yet3 can be found at all ER contact sites, it is more frequently found at the ER-mitochondria, ER-PM, and ER-LD contacts and less with ER-peroxisome contacts (Figure 1A). This is in line with findings that document a higher requirement for sterols in the former contacts^76^.

**Figure 5.**
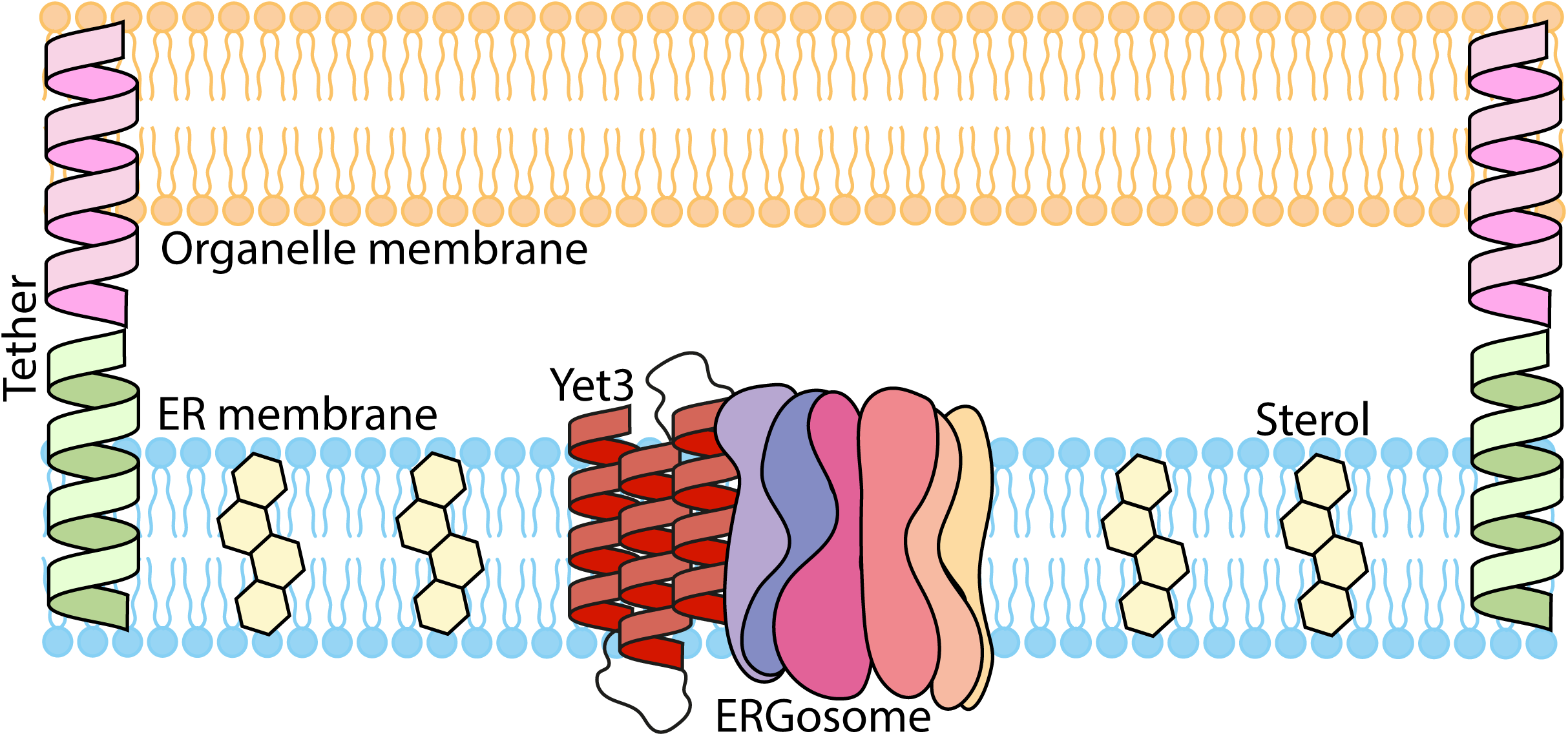
– Schematic illustration of our model for Yet3 molecular function. Yet3 accumulates at ER contact sites and recruits the post-squalene proteins in the ergosterol biosynthesis pathway to create the ERGosome. There, it increases the concentration of ergosterols, which create the sterol rich domains essential for contact formation and function.

Building ergosterols by the ERGosome requires heme, oxygen, iron, and 18 molecules of acetyl-CoA for each molecule of ergosterol (Figure S1A) ^22^. This costly sterol is therefore tightly regulated, and its formation is highly controlled. It is not surprising, therefore, that we see a “tug of war” between contact sites and the PM (the cell’s most sterol-rich membrane) for the limited amount of sterols in the cell. Indeed, we demonstrated that overexpression of Yet3 or BAP31 leads to reduced content of ergosterol/cholesterol in the PM (Figure 3B, Figure 4E). Moreover, when we compared the transcriptome profile of cells overexpressing Yet3 (Figure 3E) to the transcriptome of strains from the deletion collection^77^, we found that the closest resemblance is to Δ*erg2*— a strain deleted for an enzyme responsible for converting fecosterol to episterol in the ergosterol biosynthesis pathway (Figure S1A). This indicates that losing ergosterol biosynthesis has similar cellular consequences to depleting it from the PM by enriching sterols in contacts. This behavior can also explain the growth defect upon overexpression of Yet3 (Figure S1B). We hypothesize that elevated levels of Yet3 force the cell to direct its building blocks and energy for ergosterol synthesis, while other metabolites such as amino acids and lipids, that are required for cell duplication, are reduced (Figure S3D, Table S2, S3).

It was previously suggested that ERGosome assembly occurs at ER exit sites for vesicular trafficking^78^. This suggests that on-demand, efficient and local, sterol biosynthesis may be a widely used mechanism for creation of subdomains. Moreover, it could very well be that Yet3 may have a function in more ER subdomains, in addition to its role in ER contacts, to assemble the ERGosome, where local sterol synthesis is required. In support of this, it has been shown that Yet3 binds the SEC translocon complex which may enable translocation of specific substrates, such as GPI anchor proteins, that require high sterol and sphingolipid concentrations^70^. Another process requiring a sterol-rich subdomain is the fusion of ER tubules^79^. We found Yet3 foci co-localized with both Sey1 and Rtn1. Sey1 is the dynamin like GTPase required for fusion of tubules and Rtn1is the reticulon that maintains tubular morphology (Figure S5A). Moreover, our pull-down experiments with Yet3 also show a physical interaction with Sey1 as well as with Yop1 that works with it^80,81^ (Table S1). Indeed, looking at the ER shape, we found that overexpression of Yet3 elevates the extent of peripheral ER tubules (Figure S5B and Figure S5C). These suggest that during ER membrane shaping events, Yet3 may recruit the ERGosome to specific sites for modulating the physical properties of the lipid bilayer.

Other uses for ergosterols in the ER can be during ER stress. One of the main outcomes of ER stress is clustering of the ER stress sensor, Ire1. Ire1 clustering leads to sustained activation of the unfolded protein response (UPR) and requires upregulated sterol biosynthesis in the ER membrane^82^. It was already demonstrated that *YET3* mRNA levels^83,84^ and Yet3 protein expression^70^ are upregulated following addition of the UPR inducer dithriothreitol (DTT).

Moreover, we noticed that upon UPR induction, Yet3-GFP accumulates in specific areas on the ER (Figure S5D), which support increased Ire1 clustering overtime (Figure S5E and Figure S5F). Conversely, a dysfunctional Yet3 form (tagged on its N’) (Figure S1B) cannot support Ire1 assembly (Figure S5E). This suggests that the accumulation of Yet3 in specific areas of the ER may sequester the ERGosome to create local ergosterol enrichments required for proper Ire1 clustering.

The hypothesis that Yet3 may function in additional areas of the ER for making ergosterol-rich subdomains, can also explain our surprising observation that in many cases the depletion of *YET3* and its overexpression lead to similar phenotypes (Figure 3B, Figure S3C and Figure 3F). Indeed, depletion of *YET3* disrupts the normal accumulation of ergosterol in designated membrane areas, while overexpression of Yet3 enables the formation of the ERGosome, but recruits substantial amount of ergosterols into contacts, depleting them from other locations. In essence, the two manipulations will have similar effects on non-contact subdomains.

More mechanistically, it is not clear how Yet3 recruits the large ERGosome complex to specific domains. All ERGosome proteins harbor TMDs, and Yet3 itself possesses three predicted TMDs, which constitute approximately 50% of its protein sequence. While Yet3 and BAP31 are conserved throughout their sequence (Figure S6A), it is their TMDs that exhibit the most remarkable conservation (Figure S6B). Moreover, previous report suggests that BAP31 interacts with client membrane proteins via its TMD and may also engage with cholesterol, as demonstrated using a sterol-probe assay^85^. Interestingly, AlphaFold2 predictions^86^ suggest that as a homotrimer, Yet3 could form a hydrophobic pocket that could potentially bind ergosterol (Figure S6C). This hydrophobic channel may be important for slowing the fast sterol diffusion in the ER membrane, thus maintaining the high concentration of this molecule in the ER subdomains where Yet3 is enriched. Taken together, we hypothesize that both Yet3 and BAP31 function as scaffold proteins, facilitating the recruitment of ergosterol/cholesterol biosynthesis proteins through their TMDs. However, to fully elucidate the molecular mechanisms underlying Yet3 and BAP31 function, further structural experiments are needed.

Another intriguing question is how Yet3 itself is recruited to sites of contact to initiate local sterol accumulation. One option is that it is pulled in by interactions with VAP proteins, Scs2 and Scs22 in yeast, that also localize to all ER contacts and have been shown to interact with Yet3 (and also come up in our pull-down assays) (Table S1)^55,87^. However, in our hands, the double deletion of Scs2 and Scs22 did not abolish Yet3 accumulation at ER subdomains (Data not shown). To address this question in an unbiased approach, we imaged overexpressed Yet3-GFP on the background of the yeast knockout library (Figure S6D). This library comprises yeast strains, each carrying a targeted mutation in one specific gene^88,89^. Our preliminary results reveal that genes related to ER membrane lipid composition and LD formation, affect the localization of Yet3. Specifically, deletion of genes such as *ice2*, *loa1* and *nem1,* that under normal conditions affect the formation of di– and tri-acyl glycerol (DAG and TAG, respectively), led to a reduction in the recruitment of Yet3 to subdomains (Figure S6E). These observations suggest that the lipid milieu of the ER membrane influences the recruitment of Yet3. This may also explain why overexpressed Yet3, in contrast to other overexpressed tethers that disperse along the ER, in fact accumulates in ER subdomains. This nonstandard behavior may be the result of a feedback loop: Yet3 may accumulate itself in unique membrane micro-domains. As Yet3 accumulates, it recruits the ERGosome and increases the ergosterol concentration. Elevation of ergosterol in the contacts may encourage the assembly of more Yet3 molecules that are influenced by the ER membrane lipid composition, thus resulting in even more sterol accumulation in these subdomains and stronger Yet3 recruitment. However, to fully understand Yet3 recruitment, further comprehensive studies are needed.

Our studies focused on Yet3. However, Yet3 has two close paralogs, Yet1 and Yet2. Do they also have a role in ERGosome recruitment? A previous report^70^ and our data (Figure S4C) showed that compared to Yet1 and Yet3, Yet2 does not express constitutively, which may suggest it fulfills a different function. However, heterodimerization of Yet3 and Yet1 plays a crucial role in modulating the inositol biosynthesis pathway (Figure S2B)^55^. Moreover, deletion of *yet3* results in a reduction of Yet1 levels^70^. Surprisingly, deletion of *yet1* does not affect Yet3 expression levels but alters its distribution on the ER membrane^70^. Rather than being homogenously distributed, *yet1* deletion promotes accumulation of Yet3 in subdomains, reminiscent of the phenotype observed upon overexpression of Yet3^70^. We moreover show that deletion of *yet1,* which leads to increased recruitment of Yet3 to subdomains, also influences the distribution of ergosterol (Figure S4D). We propose therefore that the presence of Yet1 modulates Yet3 function – in its presence, heterodimerization occurs recruiting Yet3 to its role as an Opi1 regulator, while in the absence or reduced expression of Yet1, or during conditions of increased Yet3 levels – Yet3 functions independently as a scaffold for ERGosome formation. This regulatory interplay between inositol biosynthesis and ergosterol concentration underscores the multifunctional nature of Yet3. Opi1, the inhibiting factor for inositol production, also disturbs the creation of Phosphatidylethanolamine (PE) and Phosphatidylcholine (PC) by reducing the expression levels of the enzymes required to synthesize these phospholipids, Psd1, Cho2 and Opi3. This may be an efficient way to control and balance the amount of ergosterols and phospholipids in the cell. Inositol is also an essential building block for phosphatidylinositol (PI), another common phospholipid. Intriguingly, strains lacking LDs as observed in Yet3 overexpression (Figure 2A), exhibit significantly reduced PI levels even under continuous growth in the presence of inositol^90^. The reduction in overall PI content within cells overexpressing Yet3 supports our hypothesis that it serves to promote the crosstalk between both lipid metabolism pathways.

Other than the two paralogs in yeast, Yet3 also has a highly conserved human homolog, BAP31. BAP31 was reported to have roles in diverse cellular functions such as: ER stress through the UPR; lipid and glucose metabolism^41,42^; mitochondrial homeostasis; autophagy^39^; and regulation of proliferation and migration of cells^43,44^. Moreover, it is known to be involved in the activation of caspase-8 and apoptosis^36,37^. Similar to Yet3, BAP31 was shown to be a resident protein in the ER-PM contact^40^ and in MAMs, where it may regulate autophagy and apoptosis ^38,39,91^. Both pathways are known to necessitate sterol-rich domains. Although BAP31 was suggested to function as a tether protein that binds either TOM40 or FIS1^54,92^ in MAMs, this was never fully demonstrated, and a clear demonstration of a tether activity was not shown ^45^. Our results suggest that while BAP31 is indeed a resident protein in these contacts, its role may be in concentrating cholesterol at specific ER subdomains. In support of this, BAP31 interacts with SREBP1C and INSIG1, both key players in the regulation of cholesterol metabolism^93^, and with OSBP^91^, which is a lipid transfer protein that controls cholesterol transfer. BAP31 deletion induced lipogenesis and cholesterol accumulation in hepatocytes^41^ and LD accumulation in white adipose tissue^94^. Interestingly, the overexpression of both BAP31 and Yet3 resulted in reduction of LD number (Figure 2A and Figure 2B). Like Yet3, BAP31 binds proteins that are required for ER tubule shaping such as the reticulon RTN3, and the dynamin like GTPase ATL3^91^, which are also enriched in sterol rich domains. This could hint that BAP31 might also accumulate in multiple subdomains to create the optimal membrane environment for cholesterol dependent pathways.

BAP31 was first discovered due to its role in B-cell receptor maturation^35^. Since the sterol regulatory binding protein (SREBP) signaling pathway is crucial for effective antibody responses^95^, BAP31 involvement in secretion of antibodies may be through its function in cholesterol subdomain regulation. Interestingly, BAP31 selectively binds Immunoglobulin D (IgD) (and not Immunoglobulin M (IgM))^96^ and is required for optimal targeting of the Major Histocompatibility Complex (MHC) class I molecules to ER exit sites^97^. BAP31 is also involved in T cell activation and proliferation by regulating the expression of key members in the T cell receptor (TCR) signaling pathways^98^. IgD, MHC and the TCR, are all known to be enriched in sterol-rich subdomains^99^. Therefore, it could be that the reason that BAP31 knockdown increases MHC class II expression in macrophage cell surface^100^ is the opposite result of BAP31 overexpression, where we observe high concentration of cholesterol on the ER and less on the PM (Figure 4E). The connection of BAP31 to so many pathways that regulate cellular homeostasis may explain why it is associated with a variety of diseases such as different types of cancer^101–104^, neurological disorders^105,106^, metabolic syndromes^41,42^, and viral infectivity^107,108^. We suggest that BAP31 is involved in many of these diseases due to its role in regulating the cholesterol distribution in the cell.

More globally, to our knowledge, our study is the first to reveal a lipid organizing protein in contact sites. It also suggests a conserved function for BAP31 in creation of sterol-rich domains, which may explain its involvement in many cellular pathways, and surely will shed more light about its actions during development and in the course of disease.

## Acknowledgements

We thank Emma Fenech and Ofir Klein for critical reading of the manuscript. We are grateful for insightful and helpful conversations with Charles Barlowe, Robin Klemm, Tim Levine, Avner Fink, Ines Castro and Yury Bykov. We thank Michal Eisenberg-Bord for support during the initial stages of this project. We thank Emma Fenech, Mor Angel, Reut Ester Avraham, Yeynit Asraf, Hadar Meyer, Tat Cheng and Ariel Feler for helping with experimental work. We thank Yoav Peleg for cloning support and Charles Barlowe for sharing Yet3 antibody. We thank Sophie Martin from the University of Geneva and Eelco van Anken from Università Vita-Salute San Raffaele for generously sharing reagents. We thank Orly Laufman for sharing HeLa S3 cell line.

This work was supported by a collaborative grant from the Deutsche Forschungsgemeinschaft (DFG) to DR and MS (DFGRA 1028/11-1) and by a collaborative grant from the DFG to PR, RFB and MS SFB1190 (P11, P13 and P22 respectively). The robotic system of the Schuldiner lab was purchased through the kind support of the Blythe Brenden-Mann Foundation. MS is an Incumbent of the Dr. Gilbert Omenn and Martha Darling Professorial Chair in Molecular Genetics. PR is additionally supported by the DFG under Germany’s Excellence Strategy EXC 2067/1-390729940; SFB1002 (A06, PR), and the Max Planck Society. ARR is supported by the US National Institutes of Health (NIH) grants GM145350 and ES025661, a US National Science Foundation (NSF) grant 1552791, and the Georgia Institute of Technology Blanchard and Vasser-Woolley professorships. RFB is supported by the European Research Council (CoG 101088355 – cryoNERVE), and the DFG through Germany’s Excellence Strategy – EXC 2067/1-390729940.

## Materials and Methods

### Yeast strains and plasmids

The laboratory strain BY4741^109^ served as the basis for the *S. cerevisiae* strains used in this study. We performed genetic manipulations using the lithium acetate, polyethylene glycol, single-stranded DNA method^110^. The plasmids used for PCR-mediated homologous recombination have been previously documented^111,112^. We designed the primers using Primers4-Yeast^113^. Prof. Sophie Martin from the University of Geneva generously provided the pDA179-mCherry-D4H plasmid. The Ire1-mCherry plasmid was graciously given by Prof. Eelco van Anken from Università Vita-Salute San Raffaele. The plasmids and strains utilized in this study are listed in Table S6 and Table S7, respectively.

### Culturing of yeast

Yeast cells were incubated overnight at a temperature of 30°C in synthetic minimal medium, which consisted of 0.67% (wt/vol) yeast nitrogen base (YNB) with ammonium sulfate, supplemented with amino acids and 2% glucose (SD). The following antibiotics were used for selection: nourseothricin (NAT, Quimigen) at a concentration of 0.2g/l; G418 (Formedium) at a concentration of 0.5g/l; and hygromycin (HYG, Formedium) at a concentration of 0.5g/l.

Subsequently, cells were diluted and cultivated until they reached mid-logarithmic phase, characterized by an optical density (OD_600_) ranging from 0.4 to 0.9. For the different growth conditions used in this study, strains were cultured as mentioned below: 2mM DTT (Sigma); 10µg/ml Fluconazole (Sigma), medium depleted of inositol using YNB without inositol (Formedium), or medium depleted of iron, using YNB without iron.

In experiments involving the mCherry-D4H reporter for free sterols, yeast strains were diluted and incubated for approximately 3 hours at 30°C. followed by transfer to a temperature of 37°C and incubation for an additional hour prior to imaging.

### Manual fluorescence microscopy and organelle staining

Cell cultures were incubated overnight in a 96-well round-bottom plate (ThermoFisher) containing 100μl of SD media, supplemented with the necessary amino acids and/or antibiotics (as mentioned above). The cells were incubated at 30°C with slight agitation. The day after, 5μl of the overnight culture was diluted into 195μl of SD media and incubated for approximately 4 hours at 30°C with slight agitation. For the observation of the mCherry-D4H ergosterol reporter, the cultures were transferred to 37°C after approximately 3 hours. 50μl of the culture was then transferred to a 384-well glass-bottomed microscopy plate (Matrical Bioscience) coated with 0.25mg/ml Concanavalin A (ConA, Sigma). Cells in the mid-logarithmic phase were adhered to the plates by incubating at room temperature for 15 minutes. Following adherence, the cells were washed and imaged in synthetic minimal medium.

For mitochondrial staining, after adherence to the ConA, the media was replaced with synthetic minimal medium containing 50nM MitoTracker (MitoTracker Orange CMTMRos; Invitrogen) and incubated at room temperature (RT) for 10 minutes, washed once, and imaged in SD media.

For lipid droplet staining, after adherence to the ConA the media was replaced with phosphate buffered saline (PBS) containing 100µM MDH (AUTODOT™ Visualization Dye Monodansylpentane, Abgent), cells were incubated at RT for 15 minutes, washed, and imaged in PBS.

For vacuole staining, after adherence to the ConA the media was replaced with synthetic minimal medium containing 16µM FM_4-64_ (FM™ 4-64 Dye (N-(3-Triethylammoniumpropyl)-4-(6-(4-(Diethylamino) Phenyl) Hexatrienyl) Pyridinium Dibromide, Invitrogen), incubated at 30°C for 1 hour, washed, and imaged in SD media.

For vacuole area per cell quantification, images of the vacuoles with CMAC (CellTracker™ Blue CMAC (Invitrogen C2110)) dye were imaged. after adherence to the ConA the media was replaced with synthetic minimal medium containing 10µM CMAC and incubated at RT for 30 minutes, washed twice and imaged in SD media.

Imaging was performed at RT using a VisiScope Confocal Cell Explorer system, which consists of a Zeiss Yokogawa spinning disk scanning unit (CSU-W1) coupled with an inverted IX83 microscope (Olympus). Single-focal-plane and Z-stack images were acquired with a 60× oil lens (NA 1.4) and were captured using a PCO-Edge sCMOS camera, controlled by VisiView software (GFP [488 nm], RFP [561 nm], or BFP [405 nm]).

High-resolution imaging was performed at room temperature using an automated inverted fluorescence microscope system (Olympus) equipped with a spinning disk high-resolution module (Yokogawa CSU-W1 SoRa confocal scanner with double micro lenses and 50-µm pinholes). Images of cells in the 384-well plates were captured using a 60× oil lens (NA 1.42) and a Hamamatsu ORCAFlash 4.0 camera. Fluorophores were excited by a laser and images were captured in the GFP channel (excitation wavelength 488 nm, emission filter 525/50nm). All images were taken in a Z-stack, and processed using cellSens software. Images were deconvoluted using cellSens software for noise reduction and the best focal plane for presentation was selected.

Images captured using a 100X oil lens were imaged using a Hamamatsu flash orca 4.0 camera, and a CSU-W1 Confocal Scanner Unit of Yokogawa with a 50µm pinhole disk at room temperature. For images with the ergosterol sensor, the strains were maintained at 37°C during imaging. Fluorophores were excited by a laser and images were captured in two channels: GFP (excitation wavelength 488nm, emission filter 525/50nm) and mCherry (excitation wavelength 561nm, emission filter 617/73nm).

All acquired images were manually inspected and brightness adjustments were performed using ImageJ^114^. All quantifications in Figure 2A were done by ScanR Olympus soft imaging solutions version 3.2.

### Electron Tomography and Correlative Fluorescent Imaging

Pelleted yeast cells were placed in an aluminum disc with a depression of 100μm and outer diameter of 3mm (Engineering Office M. Wohlwend GmbH), then covered with a matching flat disc. The sandwiched sample was high-pressure frozen using an EM ICE high pressure-freezing device (Leica Microsystems, GmbH, Germany). Frozen samples were dehydrated in a temperature-controlled AFS2 Freeze substitution device, (Leica Microsystems) at − 90°C for 55 hours, in dry acetone containing 0.1% Uranyl Acetate (EMS, Hatfield, PA, USA). The temperature was then raised to −45°C (5°C/hour) for 9 hours followed by three acetone washes. Infiltration with Lowicryl HM20 (EMS) was carried out at increasing concentrations (10%, 25%, for 2 hours each). The temperature was then raised to −25°C (5°C/hour) and infiltration with higher concentrations of Lowicryl HM20 (50%, 75%, 2 hour each) was carried out. Finally, 100% Lowicryl HM20 was exchanged three times for every 10 hours followed by polymerization under UV light for 48 hours. The temperature was increased to 20°C (5°C/hour) and left under UV light for 48 hours. Sections of 200nm thickness were produced using an EMUC7 ultramicrotome (Leica microsystems) and were mounted on formvar coated 200 mesh copper grids.

Grids were labeled with vacuolar membrane FM_4-64_ dye (1:100, T13320, Thermo Fisher Scientific) for 15 minutes. Multi-color wide-field fluorescence imaging of the sections was performed using a VUTARA SR352 system (Bruker) in the presence of an imaging buffer composed of 5mM cysteamine, oxygen scavengers (7μM glucose oxidase and 56nM catalase) in 50mM Tris with 10mM NaCl and 10% glucose at pH 8.0. Images were recorded using 1.3 NA 60x silicon oil immersion objective (Olympus) and excitation lasers of 488nm and 561nm. Z slices of 150nm were collected in order to compensate for the curvature of the grid. Chromatic correction and alignment of the 488nm and 561nm channels was performed prior to data collection using a glass slide with tetraspeck beads (T7279, Invitrogen). Images of the channels were aligned using Vutara SRX software (Bruker).

After fluorescence mapping, grids were stained with Reynolds lead citrate for 10 minutes and transmission electron microscopy (TEM) was performed with a Tecnai TF20 Field Emission microscope (Thermo Fisher Scientific) operating at 200kV. The sample was pre-exposed to an intense flux of electrons in order to induce shrinkage of the embedded material, to prevent sample changes during tomogram acquisition. Low-magnification montaged TEM maps of the sample grids were obtained using SerialEM software^115^ The fluorescence maps were superimposed over the TEM montages using the vacuole fluorescent labeling as markers, and targets were chosen where bright puncta appeared in the cells. TEM tomograms were acquired with SerialEM, with 1nm/pixel resolution and images collected every one degree between –60 and 60 degrees. Electron tomographic datasets were reconstructed using IMOD software^116^.

High precision overlays of the fluorescence and tomographic reconstructions for specific cells was done using the vacuole labeling as a fiducial marker. For each cell, the best fluorescence Z slice was selected and overlaid with the corresponding tomographic virtual section using Adobe Photoshop. Figure 1C shows superimposed fluorescence maps overlaying virtual sections from the cellular tomographic reconstructions.

### Cryo-electron tomography

Sample vitrification: Yeast expressing either endogenous or overexpressing Yet3-GFP were cultured to OD_600_ 0.9. Glow-discharged Cryo-EM grids (R1.2/1.3, Cu 200 mesh grid, Quantifoil microtools) were mounted on a Vitrobot Mark IV (Thermo Fisher Scientific) and a drop of 3.5µL of culture was deposited on their carbon side. Excess liquid was removed by back-blotting with filter paper (Whattman 597) prior to vitrification by quick plunging into a liquid ethane/propane mixture at liquid nitrogen temperature. The grids were stored in grid boxes in liquid nitrogen until further use.

Sample thinning: Lamellae were prepared using an Aquilos 2 Cryo focused ion beam (FIB)/scanning electron microscope (SEM) (Thermo Fisher Scientific) at cryogenic temperature. A protective layer of organometallic platinum was deposited on the grid with the gas injection system (GIS) for 35 seconds. The sample was tilted to an angle of 20° and 200nm thick lamellae were prepared sequentially, starting with the Ga2+ ion beam at 30kV and 300 pA beam current for rough milling and followed by fine milling at 30kV and 50 pA. SEM imaging at 3kV and 13 pA was used to monitor the milling process.

Data acquisition: Tilt series from –46° to +64° at increments of 3° using the dose-symmetric acquisition scheme^117^ of the lamellae were acquired using a Krios G4 Cryo-transmission electron microscope (Thermo Fisher Scientific) equipped with a 300kV field emission gun, Selectris energy filter, and a Falcon 4i direct electron detector camera, at a magnification of 33000x (3.653Å/pixel) at a defocus of –5 to –7µm. The target total dose per tomogram was around 120 e-/A^2^. The camera was operated in dose-fractionation mode and between 700 to 900 EER frames were generated per tilt image. Frames were subsequently aligned using Motioncor2 and the new tilts series were reconstructed using patch tracking in IMOD 4.11^118^ and weighted back-projected to reconstruct a tomogram. Tomograms were binned to 14.61 Å/pixel and isotopically reconstructed using IsoNe^119^. Tomogram segmentation: automatic membrane segmentation was performed using MemBrain-Seg^120^. Segmentations were then manually curated and colored using Amira (Thermo Fisher Scientific). Images were produced using UCSF Chimera^121^.

### Electron Microscopy

Processing of the samples was conducted using the Tokuyasu technique (Tokuyasu, 1973). Initially, the specimens were subjected to fixation using a solution of 0.1% glutaraldehyde (EMS) and 4% paraformaldehyde (EMS) in a 0.1M cacodylate buffer (synthesized from dimethylarsinic acid sodium salt trihydrate; Sigma-Aldrich) with an addition of 5mM CaCl2 (pH 7.4; Sigma-Aldrich). This fixation process was carried out for a duration of 2 hours. Subsequently, the samples were rinsed and embedded in a 10% gelatin solution (EMS) and subjected to an additional fixation period of 24 hours at a temperature of 4°C.

Following this, the samples underwent a cryoprotection process, which involved infiltration with a 2.3M sucrose solution (J.T. Baker) for a period of 48 hours at RT, and were then frozen by immersion in liquid nitrogen. Ultrathin frozen sections, ranging from 70 to 90nm in thickness, were prepared using a Leica EM UC7 cryo-ultramicrotome. These sections were then transferred onto formvar-coated 200-mesh nickel transmission electron microscopy (EM) grids (EMS).

The grids were subsequently rinsed and embedded in a solution of 2% methyl cellulose (Sigma-Aldrich) and 0.4% uranyl acetate (EMS). The final imaging was performed using a Thermo Fisher Scientific Tecnai T12 transmission electron microscope, equipped with a bottom-mounted TVIPS TemCam-XF416 4k × 4k CMOS camera.

### Library preparation and high-throughput screening

An automated methodology was employed for the integration of specific genomic manipulations into yeast libraries, as described in previous studies^68,122^. The query strains utilized for the procedure were constructed based on a strain YMS721^123^. The handling of libraries was facilitated using a RoToR bench-top colony array instrument (Singer Instruments). In brief, the query strains were mated with library strains on rich medium plates to yield diploid cells. These cells were subsequently transferred to nitrogen starvation media for a period of seven days to induce sporulation. Haploid cells were isolated using canavanine and thialysine (Sigma-Aldrich), with the absence of leucine serving as a selection marker for MATalpha. The final library was constructed by selecting for the desired combination of manipulations. Strains representative of the final library were validated using both microscopy and check-PCR.

For the purpose of screening, libraries were imaged using a Hamamatsu flash orca 4.0 camera and a CSU-W1 Confocal Scanner Unit of Yokogawa, equipped with a 50µm pinhole disk. The ScanR Olympus soft imaging solutions acquisition 3.2 software was used for image acquisition, with images captured using a 60× air lens (NA 0.9, GFP [488 nm]). For the secondary screen aimed at hit validation, strains were imaged using a 100× oil lens (NA 0.9, GFP [488 nm]). Libraries were imaged at RT during mid-logarithmic growth phase. Manual inspection of images was performed using ImageJ software^114^.

### Drop dilution assay

Serial dilutions of the cells were cultivated on synthetic minimal medium supplemented with glucose. The cells were initially incubated overnight with the appropriate selection markers. Following this, yeast strains were back-diluted to achieve an OD_600_ of 0.2 in synthetic media and incubated for approximately 4 hours at 30°C. After undergoing at least one cell division or upon reaching mid-logarithmic phase, the strains were back-diluted again to an OD_600_ of 0.1 and subsequently diluted in 10-fold increments. 2.5µL from each dilution was then plated onto either SD agar plates or SD agar plates supplemented with 10µg/ml Fluconazole (Sigma-Aldrich), both of which contained all necessary amino acids. The plating was performed in triplicate using a multichannel pipette (Gilson). Following 3 days of growth at 30°C, images of the plates were captured using a Canon PC1591 digital camera.

### Growth assay

The growth assay was conducted in Transparent 96-well plates (Greiner) using a Spark plate reader (Tecan). The cells were incubated at a temperature of 30°C with shaking speed of 200 rpm in a Liconic incubator for 72 hours. Following a resuspension on a Bioshake 3000 plate shaker operating at 1,200 rpm, samples were measured at hourly intervals. The OD_600_ was recorded at a wavelength of 600nm.

### Western blots

Yeast strains expressing C’ GFP-tagged Yet1, Yet2, or Yet3 for figure S3C or C’ GFP-tagged Yet1 and Yet3 either endogenously expressed or overexpressed for figure S1D, were cultured at 30°C in SD complete media until they reached mid-logarithmic phase. Cell density was adjusted to an OD_600_ of 5, corresponding to approximately 20µg of protein per sample (checked by BCA). The cells were harvested by centrifugation at 3,000 x g for 3 minutes and washed with 1ml of nuclease-free water. The cell pellet was resuspended in 200µL of lysis buffer (8M urea, 50mM Tris, pH 7.5, and protease inhibitors; Merck) and lysed by vortexing with glass beads (Scientific Industries) at 4°C for 10 minutes. Subsequently, 25µL of 20% SDS was added to each sample, followed by incubation at 45°C for 15 minutes. The lysate was separated from the glass beads by piercing the bottom of the microcentrifuge tubes, placing them in 5ml tubes, and centrifuging at 4,000 x g for 10 minutes. The flow-through was transferred to a new 1.5ml microcentrifuge tube and centrifuged at 20,000 x g for 5 minutes. The supernatant was collected and mixed with 4x SDS sample buffer (Laemmli buffer) and fresh 1M DTT, followed by incubation at 45°C for 15 minutes. The protein samples were separated by SDS-PAGE on a 4–20% gradient gel (Bio-Rad) and transferred to a 0.45-µm nitrocellulose membrane (Pall Corporation) using the Trans-Blot Turbo transfer system (Bio-Rad). The membrane was blocked with SEA BLOCK buffer (Thermo Scientific; diluted 1:5 in PBS) for 1 hour at RT and incubated overnight at 4°C with primary antibodies (rabbit anti-GFP, ab290, 1:2500; Abcam and mouse anti-Actin, ab170325, 1:5000, Abcam) diluted in a 2% wt/vol BSA/PBS solution containing 0.01% NaN_3_. After washing, the membrane was probed with secondary antibodies (800CW Goat anti-Rabbit IgG, ab216773; Abcam and 680CW Goat anti-Mouse IgG, ab216776, Abcam) diluted 1:10,000 in 5% wt/vol nonfat milk/ Tris-buffered saline with 0.05% Tween 20 (TBST) for 1 hour at room temperature. The blots were washed and imaged using the LI-COR Odyssey Infrared Scanner.

### Blue Native PAGE experiments

Isolation of crude organelles was performed on yeast cells (200ml) constitutively expressing the indicated GFP tagged proteins and cultured in rich media (YP) supplemented with 2% glucose till logarithmic phase. The cells were harvested (3000 x g, 5 minutes, RT), resuspended in DTT buffer (100mM Tris, 10mM DTT) and incubated at 30°C for 15 minutes. The cells were then washed once with spheroplasting buffer (1.2M Sorbitol, 20mM KPI, pH 7.2) and incubated with spheroplasting buffer supplemented with zymolyase (6mg/g of cells) for 1 hour at 30°C, to digest the cell wall. Further steps were carried out on ice. The spheroplasts were resuspended in homogenization buffer (0.6M Sorbitol, 10mm Tris, pH 7.4, 1mM EDTA, 0.2% fatty acid-free BSA with 2mM PMSF). To obtain cell lysate, the spheroplasts were dounce homogenized. The cell debris and nuclei were removed by centrifugation (2000 x g, 10 minutes, 4°C). The supernatant containing the crude organelles were isolated by centrifugation (18,000 x g, 15 minutes, 4°C). The pellets were resuspended in SEM buffer (250mM Sucrose, 1mM EDTA, 10mM MOPS) containing 2mM PMSF and stored at –80°C.

Blue Native gels were run with the crude organelle samples (150µg) solubilized in 100µL SEM buffer supplemented with Triton X-100 with a protein to detergent ratio of 1:2. The sample was incubated for 30 minutes on ice. The supernatant containing the solubilized fraction was isolated by centrifugation (30,000 x g, 30 minutes, 4°C), and was mixed with the 10X loading dye (5% (w/v) Coomassie blue G, 500mM 6-amino-N-caproic acid, 100mM Bis-Tris, pH 7.0). The sample was loaded on a gel containing 6%-16% acrylamide gradient. The gels were run (150V, 15mA, 2 hours, 4°C) with Cathode buffer A (500mM Tricine, 150mM Bis-Tris, 0.2% Coomassie blue G, pH 7.0). The buffer was replaced with Cathode buffer B (500mM Tricine, 150mM Bis-Tris, pH 7.0) and the gels run was continued (50V, 15mA, 16 hours, 4°C). The proteins were blotted onto a polyvinylidene fluoride (PVDF) membrane and immunodecorated with a GFP antibody (Torrey Pines Biolabs Inc) in a 1:1000 dilution. Secondary antibody, Goat anti-rabbit IgG (H+L)-HRP conjugate (Bio-Rad) was applied in 1:10,000 dilution.

### Mitochondrial translation rate analysis

Wild type and overexpressed Yet3 strains were cultured overnight in YPGal. The next morning, the OD6_00_ was measured and adjusted to approximately 0.5 in YPGal. The cultures were then incubated for an additional 5 hours to ensure similar growth cycles and OD across all strains. For each reaction, 0.5 OD of culture was centrifuged at 4000 rpm for 5 minutes, washed with 500µl KPi buffer + 2% galactose, and resuspended in 500µl KPi + galactose. The cultures were then incubated for 10 minutes at 30°C in Methionine starvation medium. To block cytosolic protein synthesis, 10µl of Cycloheximide was added for 5 minutes. Subsequently, 2µl of [35S] Methionine was added for a 10-minute pulse. Then,10µl of cold Methionine was added and the samples were immediately placed on ice. The samples were then centrifuged at maximum speed for 1 minute at 4°C and frozen for further processing. The pellet was resuspended in 500µl water, mixed with 74µl of 2M NaOH and 6µl of 2-Mercaptoethanol (2-ME), and incubated on ice for 10 minutes. 80µl of 50% TCA (2M NaOH, 2-ME, TCA, acetone) was added and the mixture was incubated on ice for 10 minutes. The samples were then centrifuged at maximum speed for 20 minutes at 4°C. The pellet was finally dissolved in 50µl of 1xLoading buffer. 25µl of each sample was loaded onto a 10-18% Tris-Tricine gel. The gel was run at 25V and 250mA for 2.5 hours, stained, destained, dried, and exposed to a screen for 2 days for autoradiography.

### Immunoprecipitation and LC–MS/MS sample preparation

Approximately 5 OD_600_ of cell pellets were resuspended in 400μl of lysis buffer, which contained 150mM NaCl, 50mM Tris-HCl (pH 8.0), 5% Glycerol, 1% digitonin (Sigma), 1mM MgCl2, protease inhibitors (Merck), and benzonase (Sigma). The cell suspension was then transferred to a 2ml FastPrep™ tube (lysing matrix C, MP Biomedicals). Lysis was performed by six cycles of one minute at maximum speed on a FastPrep-24™ cell homogenizer (MP Biomedicals), with a 5 minutes cooling period on ice between each cycle. The lysates were then centrifuged at 16,000 g for 10 minutes at 4°C, and the supernatant was transferred to a new microcentrifuge tube. For the purification of Protein-GFP, the samples were incubated with 40μl of pre-washed GFP-Trap Agarose beads (Chromotek) for 1 hour at 4°C. The beads were then washed twice with 200μl of digitonin wash buffer (150mM NaCl, 50mM Tris-HCl pH 8.0, 1% digitonin) and four times with basic wash buffer (150mM NaCl, 50mM Tris-HCl pH 8.0). The beads were then incubated with 50μl of elution buffer (2M urea, 20mM Tris-HCl pH 8.0, 2mM DTT and 0.5μl trypsin (0.5μg/μl, Promega, #V5111)) per sample for 90 minutes. The eluate was then separated from the beads and collected in a new microcentrifuge tube. 50μl of alkylation buffer (2M urea, 20mM Tris-HCl pH 8.0, 50mM iodoacetamide (IAA)) was added to the beads and incubated for 10 minutes. This buffer was also separated from the beads and combined with the first eluate. Finally, the beads were washed with 50μl of urea buffer (2M urea, 20mM Tris-HCl pH 8.0) for an additional 10 minutes, and this buffer was also separated and combined with the previous mixture. All elution steps were performed at RT in the dark with shaking at 1,400 rpm. The combined eluate (150μl total volume) was incubated overnight at RT in the dark at 800 rpm. The following morning, 1μl of 0.25μg/μl trypsin was added to each sample and incubated for an additional 4 hours at RT in the dark at 800 rpm. Following digestion, peptides were desalted using Oasis HLB, μElution format (Waters, Milford, MA, USA). The samples were vacuum dried and stored in –80°C until further analysis.

### LC–MS/MS Proteomics

The mass spectrometry proteomics data have been deposited to the ProteomeXchange Consortium via the PRIDE ^124^ partner repository with the dataset identifier PXD052060 and 10.6019/PXD052060.

Liquid chromatography was used with ULC/MS grade solvents. Each sample was loaded using split-less nano-Ultra Performance Liquid Chromatography (10 kpsi nanoAcquity; Waters, Milford, MA, USA). The mobile phase was: A) H_2_O + 0.1% formic acid and B) acetonitrile + 0.1% formic acid. The peptides were then separated using a T3 HSS nano-column (75µm internal diameter, 250mm length, 1.8µm particle size; Waters) at 0.35µL/minute except for the first 13 minutes. Peptides were eluted from the column into the mass spectrometer using the following gradient: 3%B for 13 minutes at flow of 0.4µL/minute, 3% to 30%B in 42 minutes, 30% to 90%B in 5 minutes, maintained at 90% for 5 minutes and then back to initial conditions^125^.

Mass Spectrometry was performed with the nanoUPLC coupled online through a nanoESI emitter (10μm tip; New Objective; Woburn, MA, USA) to a quadrupole orbitrap mass spectrometer (Q Exactive HF, Thermo Scientific) using a FlexIon nanospray apparatus (Proxeon). Data was acquired in data dependent acquisition (DDA) mode, using a Top10 method. MS1 resolution was set to 120,000 (at 200m/z), mass range of 375-1650m/z, AGC of 3e6 and maximum injection time was set to 60 milli-seconds. MS2 resolution was set to 15,000, quadrupole isolation 1.7m/z, AGC of 1e5, dynamic exclusion of 20 seconds and maximum injection time of 60 milli-seconds^125^.

Raw data was processed with MaxQuant v2.0.1.0^126^. The data was searched with the Andromeda search engine against the SwissProt *S. cerevisiae* ATCC204508/S288c proteome database (January 2023 version, 6060 entries) in addition to the MaxQuant contaminants database. All parameters were kept as default except: Minimum peptide ratio was set to 1 and match between runs was enabled. Carbamidomethylation of C was set as a fixed modification. Oxidation of M and protein N-term acetylation were set as variable modifications. The LFQ intensities were used for further calculations using Perseus v1.6.2.3^127^. Decoy hits were filtered out, as well as proteins that were identified on the basis of a modified peptide only. The LFQ intensities were log2-transformed and only proteins that had at least 2 valid values in at least one experimental group were kept. The remaining missing values were imputed by a random low range distribution. Student’s t-tests were performed between the relevant groups to identify significant changes in protein levels.

Data analysis was done using GraphPad Prism 10.2.2.

### Metabolomics and lipidomics sample preparation

Endogenously expressed Yet3, Yet3 overexpression, and Yet3 knockout yeast strains were cultured overnight at 30°C in a synthetic minimal medium supplemented with 2% glucose and the appropriate selection marker. On the following day, 50μl of each sample was transferred to a the same medium without any selection markers, and incubated overnight at 30°C. Subsequently, the cells were diluted and grew until they reached mid-log phase, as indicated by an OD_600_ measurement. For the collection of cells, 25 OD_600_, or 10 OD_600_ specifically for free and total ergosterol level measurements, were harvested and washed twice with DDW. The cells were then flash-frozen in liquid nitrogen and stored at –80°C for future analysis.

### Metabolite extraction for polar and lipid metabolite analysis

Extraction and analysis of lipids and polar metabolites were performed as previously described ^128,129^ with some modifications: Cell pellets were extracted with 1ml of a pre-cooled (−20°C) homogenous methanol: methyl-tert-butyl-ether (MTBE) 1:3 (v/v) mixture, containing following internal standards: 0.1μg/ml of Phosphatidylcholine (17:0/17:0) (Avanti), 0.4μg/ml of Phosphatidylethanolamine (17:0/17:0, 0.15nmol/ml of Ceramide/Sphingoid Internal Standard Mixture II (Avanti, LM6005), 0.0267µg/ml d5-TG Internal Standard Mixture I (Avanti, LM6000) and 0.1μg/ml Palmitic acid-13C (Sigma, 605573). The tubes were vortexed and then sonicated for 30 minutes in an ice-cold sonication bath (taken for a brief vortex every 10 minutes). Then, UPLC-grade water (DDW): methanol (3:1, v/v) solution (0.5ml), containing internal following standards: C13 and N15 labeled amino acids standard mix (Sigma, 767964) (1:1500), was added to the tubes followed by vortex and centrifugation. The upper organic phase was transferred into a 2mL Eppendorf tube. The polar phase was re-extracted as described above, with 0.5mL of MTBE. Both organic phases were combined and dried in a Speedvac and then stored at −80°C until analysis. The lower polar phase, used for polar metabolite analysis, was treated in a similar way. For analysis, the dried lipid extracts were re-suspended in 150μl mobile phase B (see below) and centrifuged at 20800rcf (4°C for 10 minutes). Then 120µL were transferred to the HPLC vials for injection. Polar dry samples were re-suspended in 120µL methanol: DDW (50:50) and centrifuged twice to remove the debris. 70µL were transferred to the HPLC vials for injection.

Lipid extracts were analyzed by LC-MS lipidomics using a Waters ACQUITY I class UPLC system coupled to a mass spectrometer (Thermo Exactive Plus Orbitrap) which was operated in switching positive and negative ionization mode. The analysis was performed using Acquity UPLC System combined with chromatographic conditions as described^128^ with small alterations. Briefly, the chromatographic separation was performed on an ACQUITY UPLC BEH C8 column (2.1×100mm, i.d., 1.7μm) (Waters Corp., MA, USA). The mobile phase A consisted of DDW: Acetonitrile: Isopropanol 46:38:16 (v/v/v) with 1% 1M NH4Ac, 0.1% acetic acid. Mobile phase B composition is DDW: Acetonitrile: Isopropanol 1:69:30 (v/v/v) with 1% 1M NH4Ac, 0.1% acetic acid. The column was maintained at 40°C, and the flow rate of the mobile phase was 0.4ml/min. Mobile phase A was run for 1 minute at 100%, then it was gradually reduced to 25% at 12 minutes, following a decrease to 0% at 16 minutes. Then, mobile phase B was run at 100% till 21 minutes, and mobile phase A was set to 100% at 21.5 minutes. Finally, the column was equilibrated at 100% A till 25 minutes. Lipid identification and quantification was performed using LipidSearch™ software (Thermo Fisher Scientific). The validation of the putative identification of lipids was performed by comparing it to the homemade library, which contains lipids produced by various organisms, and the correlation between retention time and carbon chain length and degree of unsaturation. Relative levels of lipids were normalized to the internal standards and the protein amount in the examined samples.

LC-MS polar metabolite analysis (Metabolic profiling) was performed as described^129^ with minor modifications: Briefly, analysis was performed using Acquity I class UPLC System combined with mass spectrometer Q Exactive Plus Orbitrap™ (Thermo Fisher Scientific) which was operated in a negative ionization mode. The LC separation was done using the SeQuant Zic-pHilic (150mm × 2.1mm) with the SeQuant guard column (20mm × 2.1mm) (Merck). The Mobile phase B: acetonitrile and Mobile phase A: 20mM ammonium carbonate with 0.1% ammonia hydroxide in DDW: acetonitrile (80:20, v/v). The flow rate was kept at 200μl/minute and gradient as follow: 0-2minutes 75% of B, 14[minutes 25% of B, 18[minutes 25% of B, 19[minutes 75% of B, for 4 minutes, 23[minutes 75% of B. Data processing was done using Compound discoverer v3.3 (Thermo Fisher Scientific) software when detected compounds were identified by retention time, and fragments were verified using an in-house-generated mass spectra library.

Data analysis was done using GraphPad Prism 10.2.2.

### Targeted analysis of free ergosterol

Extraction and analysis of ergosterol were performed as previously described^130^ with some modifications: For analysis of free ergosterol the cell pellet was extracted with 400ul of a pre-cooled (−20°C) homogenous methanol: methyl-tert-butyl-ether (MTBE) 1:3 (v/v) mixture, containing epicholesterol (10uL, 1mg/mL) as internal standard. The mixture was incubated, shaking, with added zirconium beads at 30Hz, for 1.5 minutes (Retsch MM400) and then mixed at at 1,200rpm for 1 hour (Thermomixer C Eppendorf) at RT. Then, water (100uL) was added for phase separation, and the upper organic phase was collected and dried. The obtained residue was derivatized with N-trimethylsilylimidazole/ trimethylchlorosilane reagent (TMSI:TMCS, 99:1, v/v; 50uL; both from Merck) and pyridine (50uL) in a shaker at 800rpm for 1 hour at 60°C; Thermomixer C Eppendorf). The obtained solution was transferred to the GC-MS vials for injection.

Data analysis was done using GraphPad Prism 10.2.2.

### Respiration test

Yeast strains were grown on YPD plates containing NAT for a day at 30°C. The yeast colonies were transferred to liquid YPD (containing 2% glucose) and incubated overnight at 30°C with slight agitation. On the day of the experiment, cell density (OD_600_) was measured after 12 hours of growth. The cultures were diluted to an OD_600_ of 0.5 and incubated for another 2.5 hours at 30°C with slight agitation. One day before the measurement the Seahorse, XFe96/XF Pro Cell Culture microplate (Agilent) was coated with 0.1mg/ml Poly-D-Lysine (Sigma Aldrich), and was incubated with the coating solution at 4°C overnight. The yeast cells were pelleted down at 500 x g after which they were resuspended in either YPGal (containing 1% galactose) for determining basal respiration or Assay Medium (0.67% yeast nitrogen base, 2% potassium acetate, and 2% ethanol) for measuring maximal respiration^131^ to an OD_600_ of 0.1. 180μl of cell suspension was added per well to the Seahorse XFe96/XF Pro Cell Culture microplate. After seeding, the cells were incubated for 30 minutes at 30°C. The measurement was done under basal conditions and upon the addition of 40μM CCCP, and 2.5μM Antimycin A in a Seahorse XFe96 Analyzer (Agilent) at 30°C. Three cycles of mixing (for 3 minutes) and measuring (for 3 minutes) time were allotted to each condition. Data analysis was done using GraphPad Prism 10.2.2.

### Heme level measurements

Total heme measurements were performed as previously described^132^. Briefly, for all total heme measurements, ∼1×108 cells were harvested, washed in sterile ultrapure water, resuspended in 500μL of 20mM oxalic acid and stored in a closed box at 4°C overnight (16–18 hours). The next day, an equal volume (500μL) of 2M oxalic acid was added to the cell suspensions. The samples were split, with half the cell suspension transferred to a heat block set at 95°C and heated for 30 minutes and the other half of the cell suspension kept at RT for 30 minutes. All suspensions were centrifuged for 2 minutes on a table-top microfuge at 21,000g and the porphyrin fluorescence (excitation 400nm, emission 620nm) of 200μL of each sample was recorded on a Synergy Mx multi-modal plate reader using black Greiner Bio-one flat bottom fluorescence plates. Heme concentrations were calculated from a standard curve prepared by diluting 500–1500μM hemin chloride stock solutions in 0.1M NaOH into ultrapure water. In order to calculate heme concentrations, the fluorescence of the unboiled sample (taken to be the background level of protoporphyrin IX) was subtracted from the fluorescence of the boiled sample (taken to be the free base porphyrin generated upon the release of heme iron). The cellular concentration of heme was determined by dividing the moles of heme determined in this fluorescence assay by the number of cells analyzed, giving moles of heme per cell, and then converting to a cellular concentration by dividing by the volume of a yeast cell, taken to be 50fL.

Free heme measurements at steady-state was performed on cells expressing the HS1-M7A heme sensors cultured in 10mL SCE-LEU media, with or without 0.5mM succinylacetone, for ∼14–16 hours to OD_600_ ∼ 1–2. Cytosolic, nuclear and mitochondrial-targeted sensors were expressed on low copy centromeric plasmids and were driven by the *GPD* promoter (p415-GPD) ^67,132^ After culturing, cells were collected, washed in water, and resuspended in PBS solution to a density of 10 OD_600_ units per mL. 200μL of the cell suspension, corresponding to 2 OD_600_ units or 4 x 107 cells, was used to measure EGFP (excitation 488nm, emission 510nm) and mKATE2 (excitation 588nm, emission 620nm). Background autofluorescence of cells not expressing the sensors was recorded and subtracted from the EGFP and mKATE2 sensor fluorescence values. Fluorescence was recorded on a Synergy Mx multi-modal plate reader using black Greiner Bio-one flat bottom fluorescence plates.

To estimate fractional heme occupancy of the heme sensor, we used the following formula: ^133^

% Heme Occupancy = (R−R_min_) / (R_max_−R_min_) x 100

where R_min_ was the sensor ratio obtained from heme depleted cells cultured with succinylacetone, R_max_ is the ratio obtained from cells expressing HS1, a high-affinity sensor that is quantitatively saturated with heme^65^, and R is the steady-state ratio of the HS1-M7A heme sensor in a given strain.

Data analysis was performed using GraphPad Prism 10.2.2.

### RNA sample collection, extraction, and sequencing

Endogenously expressed Yet3 and overexpressed Yet3 strains were cultured overnight at 30°C in synthetic minimal medium supplemented with 2% glucose and NAT. On the following day, cells were back-diluted and cells at 2.5 OD_600_ were pelleted, washed with DDW, frozen in liquid nitrogen and stored at −80°C until further analysis. RNA was extracted using a modified protocol of the Nucleospin 96 RNA kit (Macherey-Nagel, Duren, Germany). Cells were lysed in a 96 deep-well plate by adding 450μl of lysis buffer containing 1M sorbitol (Sigma-Aldrich), 100mM EDTA, and 0.45μl lyticase (10IU/μL) and incubating at 30°C for 30 minutes. Spheroplasts were centrifuged for 10 minutes at 1,300 x g and extraction was performed as per the kit protocol substituting β-mercaptoethanol with DTT.

RNA libraries were created as follows: poly(A) RNA was selected by reverse transcription with a barcoded poly(T) primer. The barcoded DNA–RNA hybrids were pooled and fragmented by a hyperactive variant of the Tn5 transposase. Tn5 was stripped off the DNA with 0.2% SDS followed by SPRI beads cleanup, and cDNA was amplified and sequenced with Illumina Novaseq 6000 using a primer complementary to the opposite adaptor to the poly(A).

### Processing and analysis of RNA-seq data

50-bp reads of the RNA-seq of every sample were mapped to the *S. cerevisiae* genome (R64 in SGD) using bowtie2 (parameters: –very-sensitive –trim-to 40). After alignment to the genome, samples that had less than 150,000 reads were discarded from the analysis to prevent an artificial enrichment for highly expressed genes. For every sequence, we removed PCR-duplicates using UMIs and umitools. For each gene, we summed all the unique reads aligned to 400 bp upstream its 3′ end to 200-bp downstream in order to get the total expression of that gene. The number of reads for each sample was normalized to 10^6^. Expression fold change was calculated between the log2-normalized means of the different samples and the p-value calculated based on expression values in individual repeats (n=3).

Data analysis was done using GraphPad Prism 10.2.2.

### Mammalian cell culture

Human HeLa S3 cells (graciously shared by Orly Laufman, Weizmann Institute of Science) were maintained in DMEM/F-12 medium (Gibco, USA) supplemented with 10% fetal bovine serum (FBS, Gibco, USA) and 1% Penicillin-Streptomycin-Neomycin (Biological Industries, Israel).

### Mammalian Plasmids and Transfections

HeLa S3 cells were transfected with plasmids expressing the pLenti-C-mGFP-P2A-Puro Lentiviral Gene Expression Vector (Origene) or BAP31 (BCAP3) Human Tagged Lenti ORF Clone plasmids (Origene) (1µg/ml) for 4 hours using Lipofectamine 2000 (Invitrogen, USA) and then were grown in DMEM/F-12 medium overnight.

### Imaging and staining of Mammalian Cells

Cells were cultured on coverslips placed in a 12-well plate and subsequently fixed using 4% paraformaldehyde (PFA) for a duration of 20 minutes. The nuclei were stained with Hoechst H33342 (1:1000 dilution, Sigma, USA) for approximately 10 minutes, while LDs were stained with BODIPY 558/568 (1:2000 dilution, Invitrogen, D3835) for ∼45 minutes at RT. The coverslips were then mounted using Immu-Mount mounting medium (Thermofisher scientific, USA).

For the purpose of cholesterol staining, cells were grown on a 12-well plate and fixed in 4% PFA for 15 minutes at RT. Following fixation, the cells were washed thrice with PBS. The fixed cells were then incubated with freshly prepared filipin III (F9765; Sigma-Aldrich) at a concentration of 50 µg/ml in PBS for ∼1.5 hours at RT in the dark. After three additional washes with PBS, the cells were mounted using Immu-Mount mounting medium.

Image acquisition was performed using an Andor Dragonfly 505 confocal spinning disk system, operated via Fusion software. The system is integrated with a Leica Dmi8 inverted microscope, and images were captured using a Plan Apo 63x (1.40 N.A.) oil immersion lens. All images were subsequently edited using the Fiji image processing package^114^.

### Protein modelling and analysis

Yet3 and BAP31 sequences were retrieved from UniProt (PMC9825514) and monomeric or homotrimeric structures were obtain with ColabFold’s (PMC9184281) implementation of AlphaFold2 (PMC8371605) (AlphaFold2_advanced_v2). Structures were visualized in ChimeraX (PMC10588335) software. Structural based alignments from ChimeraX were rendered in ESPript3 (PMC4086106). Hydrophobic cavities were calculated with MOLE (PMC6030847).

## Figure Legends

**Figure S1.** – Overexpression of Yet3 tagged on its C’ has the same growth rate as untagged Yet3, and accumulates at ER subdomains. A. Illustration of the ergosterol biosynthesis pathway. The scheme contains all the enzymes and metabolites required for ergosterol synthesis, divided to the pre-squalene and post-squalene steps. Fluconazole, an inhibitor of Erg11, reduces ergosterol amounts in the cell. B. A growth assay of strains overexpressing (OE) Yet3 fused to GFP on either its C’ or N’. While overexpressed, N’ tagged Yet3 shows the same growth rate as control (BY4741 strain) and Yet3 knockout (KO), the C’ tagged overexpressed Yet3 has a similar growth rate to an untagged overexpressed Yet3 strain, demonstrating that tagging on this terminus preserves the function of the protein. Strains were grown overnight in synthetic minimal medium, back diluted to an OD_600_ of ∼0.01 and monitored for growth over 48 hours. C. Yet3-GFP overexpression shows high protein concentration at specific subdomains of the ER. One of the Yet3 paralogs in yeast, Yet1-GFP, is homogenously distributed across the ER both when endogenously expressed and when overexpressed. GFP intensity between the control and overexpressed strains was adjusted for brightness. Images were taken with a 100x oil lens. Scale bar, 5µm. D. A strong promoter for overexpressing (OE) Yet1 and Yet3 results in higher protein levels. Western blot analysis of strains with Yet1 and Yet3, either endogenously expressed or overexpressed from a *TEF* promoter, all C’ tagged with GFP. Immunoblotting was performed with antibodies against GFP and Actin, as a loading control.

**Figure S2.** – Yet3 overexpression affects mitochondrial morphology. A. Mitochondria are enlarged in overexpressed (OE) Yet3 compared to a control strain. EM images of yeast show the mitochondrial size in different Yet3 expression levels. M, mitochondrion. Scale bar, 200nm. B. Schematic illustration of the inositol biosynthesis regulation in yeast. In an inositol-rich environment, Opi1 inhibits the transcription activators Ino2/Ino4 and prevents expression of Ino1 and other phospholipid synthesis related-proteins. Depletion of inositol from the media results in sequestering Opi1 to the nuclear-ER, where it binds the Yet1-Yet3 heterodimeric complex, together with Scs2.

**Figure S3.** – Overexpression of Yet3 rewires cellular metabolism. A. Free ergosterol measurements show lower concentrations of ergosterols in overexpression (OE) of Yet3 compared to the control strain. GC-MS lipid profile of sterols extracted from control, overexpressed Yet3, and Yet3 knockout (KO) strains in mid-logarithmic growth phase were analyzed. P-values were calculated from the three replicates of each strain using a one-way ANOVA test. *p ≤ 0.05, **p<0.01, NS=non-significant. B. Long-chain sphingolipids (SPH d22:2) are more abundant in strains overexpressing (OE) Yet3, that also have lower ergosterol levels (Figure S3A). Extraction and analysis of lipids by lipidomic analysis was done on independent triplicates from each strain. One-way ANOVA was tested for significance. *p ≤ 0.05, NS=non-significant. C. Gap1 and Agp1, two amino acid permeases that require ergosterol rich regions for trafficking to the plasma membrane, are mis-localized due to altered expression of Yet3. Both proteins were tagged with GFP on their C’. Strains were imaged using a 60x oil lens. Scale bar, 5µm. D. Levels of various amino acids in overexpression (OE) of Yet3 compared to control. Triplicates from each sample were analyzed using polar metabolite profiling. Presented are only amino acids with significant changes as tested using two-way ANOVA. *p<0.05, **p<0.01, ****p ≤ 0.0001. E. The transcription factor Gcn4 enters the nucleus when Yet3 is overexpressed (OE). Gcn4 was tagged with GFP on its C’. Images were taken with a 60X oil lens during their logarithmic growth phase. Scale bar, 5µm. F. Overexpression (OE) of Yet3 reduces the translation rate of mitochondrial-encoded proteins. Translation rate was tested using a mitochondrial translation assay. Three replicates were performed and significance was tested by a two-tailed unpaired t-test, **p<0.01. G. Different expression levels of Yet3 do not affect basal Oxygen Consumption Rate (OCR) in non-fermentable carbon source. Yeast strains were grown on glucose and then transferred to ethanol-based buffer for Seahorse measurements. Overexpression (OE) of Yet3 did not show any significant differences compared to the control strain. Knockout (KO) of Yet3 OCR had a small difference compared to control OCR. Three biological repeats and two technical repeats were performed. *p<0.05, NS=non-significant. H. All intermediate metabolites of glycolysis are reduced in overexpression (OE) of Yet3. The exit metabolite in glycolysis, Pyruvic acid, is increased in Yet3 overexpression. Presented are only metabolites with significant changes. Triplicates from control and overexpressed strains were analyzed using polar metabolite profiling. Significant changes were tested using two-way ANOVA. **p<0.01, ***p ≤ 0.001, ****p ≤ 0.0001. I. TCA cycle intermediate metabolites are reduced in overexpression (OE) of Yet3. Citrate, the entry point for the TCA cycle, is increased in Yet3 overexpression. Presented are only metabolites with significant changes. Triplicates from control and overexpression strains were analyzed using polar metabolite profiling. Significant changes were tested using two-way ANOVA. ****p ≤ 0.0001.

**Figure S4.** – Yet3 assembles the post-squalene biosynthesis enzymes at ER contact sites, independently from its paralog Yet1. A. Overexpression (OE) of Yet3 changes the positioning of proteins involved in the post-squalene biosynthesis pathway, directing them to subdomains on the ER membrane. In strains of Yet3-GFP expressed at normal levels, post-squalene proteins, which were tagged with mCherry at their N’ and under control of a constitutive promoter, were evenly distributed. Overexpressed Yet3 recruits these proteins into specific ER subdomains that aligned with Yet3 puncta. The fact that the pre-squalene protein, mCherry-Hmg2, did not change its localization shows that this is not due to a general reorganization of ER proteins. The images were captured using a 100x oil lens. Scale bar, 5µm. B. Potential high molecular weight complexes observed by blue native gel analysis. Two potential complexes of the ERGosome together with Yet3 are shown migrating at ∼720kDa and ∼480kDa, emphasized by red triangles. Yet3 is overexpressed and tagged with GFP on its C’ while the ergosterol proteins are constitutively expressed and tagged with GFP on their N’. Crude preparations enriched for mitochondria and the ER were analysed and probed with anti-GFP antibody. C. Endogenous Yet3 is more abundant than its two paralogs Yet1 and Yet2. Western blot analysis of three different strains, Yet1, Yet2 and Yet3, all tagged with GFP on their C’ and under regulation of their native promoter. Immunoblotting was performed with antibodies against GFP and Actin as a loading control. D. Yet1 expression levels affect the presence of Yet3 puncta amount and size, and change the free ergosterol distribution in the cell. Yet1 knockout (KO) decreases the signal of overexpressed (OE) Yet3 in the ER and increases its punctate distribution. Under these conditions, the free ergosterol is more concentrated inside the cell. However, Yet1 overexpression causes Yet3 to redistribute homogeneously on the ER surface and restores ergosterol distribution on the PM. Yet3 was, in both strains, tagged with GFP on its C’. Free ergosterols were tracked by mCherry-D4H reporter. White arrows represent co-localization between the mCherry-D4H and Yet3 foci. All samples were imaged with a 100x oil lens. Scale bar, 5µm. E. Reduced amount of cellular ergosterols lead to accumulation of Yet3 in ER subdomains together with the ergosterol biosynthesis pathway proteins. Addition of 10ug/ml Fluconazole, caused endogenous Yet3-GFP signal that is usually distributed homogeneously around the ER, to accumulate in ER subdomains. Different constitutively expressed ergosterol biosynthesis proteins tagged with mCherry on their N’ moved from their even distribution on the ER membrane or the LD, to ER subdomains too when treated with Fluconazole. White arrows highlight cases of co-localization with Yet3. As a control, the localization of Hmg1, a pre-squalene pathway enzyme, was not affected by Fluconazole. Images of the strains were taken with a 100x oil lens. Scale bar, 5µm. F. Iron depletion from the media enhance overexpressed (OE) Yet3 puncta shape and induced endogenous Yet3 accumulation in ER subdomains. Yet3-GFP strains, either endogenously expressed or overexpressed, were grown under iron-free conditions. In these conditions, Yet3 shows enhanced localization to subdomains unlike the control ER protein Sec63 tagged with GFP on its C’ remained that remained distributed along the entire ER membrane. White arrows highlight Yet3 foci formation in lower concentration of iron. The images were captured using a 60x oil lens Scale bar, 5µm.

**Figure S5.** – Yet3 may recruit the ERGosome to additional ER subdomains that require ergosterol rich areas to function. A. Sey1 and Rtn1 accumulations co-localize with Yet3 at ER subdomains. Yet3 was overexpressed (OE) and C’ tagged with GFP. Sey1 and Rtn1 were constitutively expressed and tagged with mCherry on their N’. White arrows indicate the co-localization of Yet3 puncta with Sey1 and Rtn1 accumulation. Images were taken in 100x magnification. Scale bar, 5µm. B. Overexpression (OE) of Yet3 shows a higher extent of ER tubules at the cell periphery. The ER was marked by overexpression of Emc6 tagged on its N’. Both images were taken in the same Z-plane (that highlights peripheral ER tubules) using a 60x oil lens SORA. Scale bar, 5µm. C. Increased number of ER tubules in overexpressed (OE) Yet3 as visualized using cryo-ET. Tomographic slices (left) and corresponding segmentations of cryo-FIB (right) milled strains expressing endogenous (top) or overexpressing (bottom) Yet3-GFP. Scale bars, 200nm. D. Endogenous Yet3-GFP accumulates in specific ER subdomains upon DTT treatment. Yet3 tagged with GFP on its C’ was grown overnight in synthetic minimal media and back diluted to 2mM DTT for 4 hours before imaging. White arrows show Yet3 concentration in the ER membrane. Images were taken with a 60x oil lens. Scale bar, 5µm. E. The functional form of Yet3 is necessary for clustering Ire1 in response to ER stress induced by DTT (2mM, 4 hours). Overexpression (OE) of Yet3 tagged on its C’ (the functional form, Figure S1B) enables Ire1 clustering, while tagging on its N’ (non-functional form) does not. Ire1-mCherry was visualized using a plasmid. Scale bar, 5µm. F. Quantification of Ire1 clustering upon UPR induction by DTT treatment. Overexpression (OE) of Yet3 increased the percentage of cells with visible Ire1 clusters compared to control cells after DTT induction (2mM for 2 hours). Cells were counted manually using Image J.

**Figure S6.** – Yet3 concentration in subdomains are dependent on proteins that change the ER membrane lipid composition. A. Comparison of the Yet3 and BAP31 amino acid sequence and secondary structure showing high conservation of the transmembrane domains (TMDs). Each α represents one alpha helix of the proteins, starting from the N’ until the C’. Red triangles are identical amino acids situated in the same place for both Yet3 and BAP31. White triangles represent amino acids with the same charge. B. Yet3 and BAP31 TMDs are highly conserved according to AlphaFold2 prediction. A structural based sequence alignment of Yet3 (yellow) and BAP31 (blue) monomers. The N’ of both Yet3 and BAP31 contain the three TMDs and the C’ faces the cytosol. C. A model of a Yet3 homotrimer predicted using AlphaFold2. Three Yet3 molecules with three membrane spanning domains each, create a hydrophobic cavity (seen in purple). D. Schematic of the high content screen for finding genes that affect Yet3’s capacity to localize to subdomains. Overexpression (OE) of Yet3 tagged with GFP on its C’ was integrated to a deletion/hypomorphic allele library. In this library, every colony harbors a loss-of-function mutant in each of the ∼6,000 yeast genes. All strains were imaged by automated microscopy, followed by manual inspection to identify proteins that are either required for, or suppress, the subdomain localization. E. Results of the screen to uncover which mutated genes affected Yet3 accumulation at a specific subdomain of the ER. Deletion of either *ice2*, *loa1* or *nem1*, all proteins related to the ER lipid composition and LD formation, reduces Yet3 subdomain association. For a full list of hits see Supplementary Table S8. Scale bar, 5µm.

## References

1. Zung, N., & Schuldiner, M. (2020). New horizons in mitochondrial contact site research. In Biological Chemistry (Vol. 401, Issue 6). 10.1515/hsz-2020-0133.

2. Eisenberg-Bord, M., Shai, N., Schuldiner, M., & Bohnert, M. (2016). A Tether Is a Tether Is a Tether: Tethering at Membrane Contact Sites. In Developmental Cell (Vol. 39, Issue 4). 10.1016/j.devcel.2016.10.022.

3. Scorrano, L., De Matteis, M.A., Emr, S., Giordano, F., Hajnóczky, G., Kornmann, B., Lackner, L.L., Levine, T.P., Pellegrini, L., Reinisch, K., et al. (2019). Coming together to define membrane contact sites. Nat Commun 10. 10.1038/s41467-019-09253-3.

4. Shai, N., Yifrach, E., Van Roermund, C.W.T., Cohen, N., Bibi, C., Ijlst, L., Cavellini, L., Meurisse, J., Schuster, R., Zada, L., et al. (2018). Systematic mapping of contact sites reveals tethers and a function for the peroxisome-mitochondria contact. Nat Commun 9. 10.1038/s41467-018-03957-8.

5. Bernhard, W., and Rouiller, C. (1956). Close topographical relationship between mitochondria and ergastoplasm of liver cells in a definite phase of cellular activity. J Biophys Biochem Cytol 2, 73–78. 10.1083/JCB.2.4.73.

6. Phillips, M.J., and Voeltz, G.K. (2015). Structure and function of ER membrane contact sites with other organelles. Nature Reviews Molecular Cell Biology 2015 17:2 17, 69–82. 10.1038/nrm.2015.8.

7. Prinz, W. A., Toulmay, A., & Balla, T. (2020). The functional universe of membrane contact sites. In Nature Reviews Molecular Cell Biology (Vol. 21, Issue 1). 10.1038/s41580-019-0180-9

8. Castro, I.G., Shortill, S.P., Dziurdzik, S.K., Cadou, A., Ganesan, S., Valenti, R., David, Y., Davey, M., Mattes, C., Thomas, F.B., et al. (2022). Systematic analysis of membrane contact sites in Saccharomyces cerevisiae uncovers modulators of cellular lipid distribution. Elife 11. 10.7554/ELIFE.74602.

9. Eisenberg-Bord, M., Zung, N., Collado, J., Drwesh, L., Fenech, E.J., Fadel, A., Dezorella, N., Bykov, Y.S., Rapaport, D., Fernandez-Busnadiego, R., et al. (2021). Cnm1 mediates nucleus-mitochondria contact site formation in response to phospholipid levels. J Cell Biol 220. 10.1083/JCB.202104100.

10. Elbaz-Alon, Y., Rosenfeld-Gur, E., Shinder, V., Futerman, A.H., Geiger, T., and Schuldiner, M. (2014). A dynamic interface between vacuoles and mitochondria in yeast. Dev Cell 30, 95–102. 10.1016/j.devcel.2014.06.007.

11. Cohen, Y., Klug, Y.A., Dimitrov, L., Erez, Z., Chuartzman, S.G., Elinger, D., Yofe, I., Soliman, K., Gärtner, J., Thoms, S., et al. (2014). Peroxisomes are juxtaposed to strategic sites on mitochondria. Mol Biosyst 10. 10.1039/c4mb00001c.

12. Ardail, D., Popa, I., Bodennec, J., Louisot, P., Schmitt, D., and Portoukalian, J. (2003). The mitochondria-associated endoplasmic-reticulum subcompartment (MAM fraction) of rat liver contains highly active sphingolipid-specific glycosyltransferases. Biochemical Journal 371. 10.1042/BJ20021834.

13. Fujimoto, M., Hayashi, T., and Su, T.P. (2012). The role of cholesterol in the association of endoplasmic reticulum membranes with mitochondria. Biochem Biophys Res Commun 417. 10.1016/j.bbrc.2011.12.022.

14. Hayashi, T., and Fujimoto, M. (2010). Detergent-resistant microdomains determine the localization of σ-1 receptors to the endoplasmic reticulum-mitochondria junction. Mol Pharmacol 77. 10.1124/mol.109.062539.

15. Pani, B., Hwei, L.O., Liu, X., Rauser, K., Ambudkar, I.S., and Singh, B.B. (2008). Lipid rafts determine clustering of STIM1 in endoplasmic reticulum-plasma membrane junctions and regulation of store-operated Ca2+ entry (SOCE). Journal of Biological Chemistry 283. 10.1074/jbc.M800107200.

16. Garofalo, T., Matarrese, P., Manganelli, V., Marconi, M., Tinari, A., Gambardella, L., Faggioni, A., Misasi, R., Sorice, M., and Malorni, W. (2016). Evidence for the involvement of lipid rafts localized at the ER-mitochondria associated membranes in autophagosome formation. Autophagy 12. 10.1080/15548627.2016.1160971.

17. Sano, R., Annunziata, I., Patterson, A., Moshiach, S., Gomero, E., Opferman, J., Forte, M., and d’Azzo, A. (2009). GM1-Ganglioside Accumulation at the Mitochondria-Associated ER Membranes Links ER Stress to Ca2+-Dependent Mitochondrial Apoptosis. Mol Cell 36. 10.1016/j.molcel.2009.10.021.

18. Manganelli, V., Matarrese, P., Antonioli, M., Gambardella, L., Vescovo, T., Gretzmeier, C., Longo, A., Capozzi, A., Recalchi, S., Riitano, G., et al. (2021). Raft-like lipid microdomains drive autophagy initiation via AMBRA1-ERLIN1 molecular association within MAMs. Autophagy 17. 10.1080/15548627.2020.1834207.

19. Hayashi, T., and Su, T.P. (2010). Cholesterol at the endoplasmic reticulum: Roles of the sigma-1 receptor chaperone and implications thereof in human diseases. Subcell Biochem 51. 10.1007/978-90-481-8622-8_13.

20. Montesinos, J., Pera, M., Larrea, D., Guardia-Laguarta, C., Agrawal, R.R., Velasco, K.R., Yun, T.D., Stavrovskaya, I.G., Xu, Y., Koo, S.Y., et al. (2020). The Alzheimer’s disease-associated C99 fragment of APP regulates cellular cholesterol trafficking. EMBO J 39. 10.15252/embj.2019103791.

21. Area-Gomez, E., Del Carmen Lara Castillo, M., Tambini, M.D., Guardia-Laguarta, C., De Groof, A.J.C., Madra, M., Ikenouchi, J., Umeda, M., Bird, T.D., Sturley, S.L., et al. (2012). Upregulated function of mitochondria-associated ER membranes in Alzheimer disease. EMBO Journal 31. 10.1038/emboj.2012.202.

22. Jordá, T., & Puig, S. (2020). Regulation of ergosterol biosynthesis in saccharomyces cerevisiae. In Genes (Vol. 11, Issue 7). 10.3390/genes11070795

23. Sokolov, S. S., Trushina, N. I., Severin, F. F., & Knorre, D. A. (2019). Ergosterol Turnover in Yeast: An Interplay between Biosynthesis and Transport. In Biochemistry (Moscow) (Vol. 84, Issue 4). 10.1134/S0006297919040023

24. Zheng Koh, D. H., & Saheki, Y. (2021). Regulation of Plasma Membrane Sterol Homeostasis by Nonvesicular Lipid Transport. In Contact (Vol. 4). 10.1177/25152564211042451

25. Gu, Y., Jiao, X., Ye, L., & Yu, H. (2021). Metabolic engineering strategies for de novo biosynthesis of sterols and steroids in yeast. In Bioresources and Bioprocessing (Vol. 8, Issue 1). 10.1186/s40643-021-00460-9

26. Olzmann, J. A., & Carvalho, P. (2019). Dynamics and functions of lipid droplets. In Nature Reviews Molecular Cell Biology (Vol. 20, Issue 3). 10.1038/s41580-018-0085-z.,

27. Mesmin, B., & Maxfield, F. R. (2009). Intracellular sterol dynamics. In Biochimica et Biophysica Acta – Molecular and Cell Biology of Lipids (Vol. 1791, Issue 7). 10.1016/j.bbalip.2009.03.002

28. Mo, C., and Bard, M. (2005). A systematic study of yeast sterol biosynthetic protein-protein interactions using the split-ubiquitin system. Biochim Biophys Acta Mol Cell Biol Lipids 1737. 10.1016/j.bbalip.2005.11.002.

29. Kumar, N., Leonzino, M., Hancock-Cerutti, W., Horenkamp, F.A., Li, P.Q., Lees, J.A., Wheeler, H., Reinisch, K.M., and De Camilli, P. (2018). VPS13A and VPS13C are lipid transport proteins differentially localized at ER contact sites. Journal of Cell Biology 217. 10.1083/JCB.201807019.

30. Elbaz-Alon, Y., Eisenberg-Bord, M., Shinder, V., Stiller, S.B., Shimoni, E., Wiedemann, N., Geiger, T., and Schuldiner, M. (2015). Lam6 Regulates the Extent of Contacts between Organelles. Cell Rep 12. 10.1016/j.celrep.2015.06.022.

31. Costello, J.L., Castro, I.G., Hacker, C., Schrader, T.A., Metz, J., Zeuschner, D., Azadi, A.S., Godinho, L.F., Costina, V., Findeisen, P., et al. (2017). ACBD5 and VAPB mediate membrane associations between peroxisomes and the ER. Journal of Cell Biology 216. 10.1083/jcb.201607055.

32. Loewen, C.J.R., and Levine, T.P. (2005). A highly conserved binding site in vesicle-associated membrane protein-associated protein (VAP) for the FFAT motif of lipid-binding proteins. Journal of Biological Chemistry 280. 10.1074/jbc.M500147200.

33. Murley, A., Sarsam, R.D., Toulmay, A., Yamada, J., Prinz, W.A., and Nunnari, J. (2015). Ltc1 is an ER-localized sterol transporter and a component of ER-mitochondria and ER-vacuole contacts. Journal of Cell Biology 209. 10.1083/jcb.201502033.

34. Toikkanen, J.H., Fatal, N., Hildén, P., Makarow, M., and Kuismanen, E. (2006). YET1, YET2 and YET3 of Saccharomyces cerevisiae encode BAP31 homologs with partially overlapping functions. Journal of Biological Sciences 6, 446–456. 10.3923/JBS.2006.446.456.

35. Kim, K.M., Adachi, T., Peter, J.N., Terashima, M., Lamers, M.C., Köhler, G., and Reth, M. (1994). Two new proteins preferentially associated with membrane immunoglobulin D. EMBO J 13, 3793. 10.1002/J.1460-2075.1994.TB06690.X.

36. Ng, F.W.H., Nguyen, M., Kwan, T., Branton, P.E., Nicholson, D.W., Cromlish, J.A., and Shore, G.C. (1997). p28 Bap31, a Bcl-2/Bcl-XL– and procaspase-8-associated protein in the endoplasmic reticulum. J Cell Biol 139, 327–338. 10.1083/JCB.139.2.327.

37. Breckenridge, D. G., Nguyen, M., Kuppig, S., Reth, M., & Shore, G. C. (2002). The procaspase-8 isoform, procaspase-8L, recruited to the BAP31 complex at the endoplasmic reticulum. Proceedings of the National Academy of Sciences of the United States of America, 99(7), 4331–4336. 10.1073/PNAS.072088099

38. Iwasawa, R., Mahul-Mellier, A.L., Datler, C., Pazarentzos, E., and Grimm, S. (2011). Fis1 and Bap31 bridge the mitochondria-ER interface to establish a platform for apoptosis induction. EMBO Journal 30. 10.1038/emboj.2010.346.

39. Namba, T. (2019). BAP31 regulates mitochondrial function via interaction with Tom40 within ER-mitochondria contact sites. Sci Adv 5. 10.1126/SCIADV.AAW1386.

40. Jing, J., He, L., Sun, A., Quintana, A., Ding, Y., Ma, G., Tan, P., Liang, X., Zheng, X., Chen, L., et al. (2015). Proteomic mapping of ER-PM junctions identifies STIMATE as a regulator of Ca 2+ influx. Nat Cell Biol 17. 10.1038/ncb3234.

41. Xu, J.L., Li, L.Y., Wang, Y.Q., Li, Y.Q., Shan, M., Sun, S.Z., Yu, Y., and Wang, B. (2018). Hepatocyte-specific deletion of BAP31 promotes SREBP1C activation, promotes hepatic lipid accumulation, and worsens IR in mice. J Lipid Res 59, 35. 10.1194/JLR.M077016.

42. Wu, Z., Yang, F., Jiang, S., Sun, X., and Xu, J. (2018). Induction of Liver Steatosis in BAP31-Deficient Mice Burdened with Tunicamycin-Induced Endoplasmic Reticulum Stress. Int J Mol Sci 19. 10.3390/IJMS19082291.

43. Dang, E., Yang, S., Song, C., Jiang, D., Li, Z., Fan, W., Sun, Y., Tao, L., Wang, J., Liu, T., et al. (2018). BAP31, a newly defined cancer/testis antigen, regulates proliferation, migration, and invasion to promote cervical cancer progression. Cell Death & Disease 2018 9:8 9, 1–15. 10.1038/s41419-018-0824-2.

44. Kim, W.T., Choi, H.S., Lee, H.M., Jang, Y.J., and Ryu, C.J. (2014). B-cell receptor-associated protein 31 regulates human embryonic stem cell adhesion, stemness, and survival via control of epithelial cell adhesion molecule. Stem Cells 32, 2626–2641. 10.1002/STEM.1765.

45. Eisenberg-Bord, M., Shai, N., Schuldiner, M., & Bohnert, M. (2016). A Tether Is a Tether Is a Tether: Tethering at Membrane Contact Sites. In Developmental Cell (Vol. 39, Issue 4). 10.1016/j.devcel.2016.10.022

46. Shai, N., Yifrach, E., Van Roermund, C.W.T., Cohen, N., Bibi, C., Ijlst, L., Cavellini, L., Meurisse, J., Schuster, R., Zada, L., et al. (2018). Systematic mapping of contact sites reveals tethers and a function for the peroxisome-mitochondria contact. Nat Commun 9. 10.1038/s41467-018-03957-8.

47. Gatta, A.T., Wong, L.H., Sere, Y.Y., Calderón-Noreña, D.M., Cockcroft, S., Menon, A.K., and Levine, T.P. (2015). A new family of StART domain proteins at membrane contact sites has a role in ER-PM sterol transport. Elife 4. 10.7554/eLife.07253.

48. Manford, A.G., Stefan, C.J., Yuan, H.L., MacGurn, J.A., and Emr, S.D. (2012). ER-to-Plasma Membrane Tethering Proteins Regulate Cell Signaling and ER Morphology. Dev Cell 23. 10.1016/j.devcel.2012.11.004.

49. Kornmann, B., Kornmann, B., Currie, E., Currie, E., Collins, S.R., Collins, S.R., Schuldiner, M., Schuldiner, M., Nunnari, J., Nunnari, J., et al. (2009). An ER-Mitochondria Tethering Complex Revealed by a Synthetic Biology Screen. Science (1979) 325.

50. Pan, X., Roberts, P., Chen, Y., Kvam, E., Shulga, N., Huang, K., Lemmon, S., and Goldfarb, D.S. (2000). Nucleus-vacuole junctions in Saccharomyces cerevisiae are formed through the direct interaction of Vac8p with Nvj1p. Mol Biol Cell 11. 10.1091/mbc.11.7.2445.

51. Toulmay, A., and Prinz, W.A. (2012). A conserved membrane-binding domain targets proteins to organelle contact sites. J Cell Sci 125. 10.1242/jcs.085118.

52. Lackner, L.L., Ping, H., Graef, M., Murley, A., and Nunnari, J. (2013). Endoplasmic reticulum-associated mitochondria-cortex tether functions in the distribution and inheritance of mitochondria. Proc Natl Acad Sci U S A 110. 10.1073/pnas.1215232110.

53. Geiger, R., Andritschke, D., Friebe, S., Herzog, F., Luisoni, S., Heger, T., and Helenius, A. (2011). BAP31 and BiP are essential for dislocation of SV40 from the endoplasmic reticulum to the cytosol. Nat Cell Biol 13, 1305–1314. 10.1038/NCB2339.

54. Namba, T. (2019). BAP31 regulates mitochondrial function via interaction with Tom40 within ER-mitochondria contact sites. Sci Adv 5. 10.1126/SCIADV.AAW1386.

55. Wilson, J.D., Thompson, S.L., and Barlowe, C. (2011). Yet1p-Yet3p interacts with Scs2p-Opi1p to regulate ER localization of the Opi1p repressor. 10.1091/mbc.E10-07-0559.

56. Wagner, C., Blank, M., Strohmann, B., & Schüller, H. J. (1999). Overproduction of the Opi1 repressor inhibits transcriptional activation of structural genes required for phospholipid biosynthesis in the yeast Saccharomyces cerevisiae. Yeast, 15(10 A). 10.1002/(SICI)1097-0061(199907)15:10A<843::AID-YEA424>3.0.CO;2-M

57. Marek, M., Vincenzetti, V., and Martin, S.G. (2020). Sterol biosensor reveals LAM-family ltc1-dependent sterol flow to endosomes upon arp2/3 inhibition. Journal of Cell Biology 219. 10.1083/JCB.202001147.

58. Berkow, E. L., & Lockhart, S. R. (2017). Fluconazole resistance in Candida species: A current perspective. In Infection and Drug Resistance (Vol. 10). 10.2147/IDR.S118892

59. Kim, Y., Mavodza, G., Senkal, C.E., and Burd, C.G. (2023). Cholesterol-dependent homeostatic regulation of very long chain sphingolipid synthesis. Journal of Cell Biology 222. 10.1083/jcb.202308055.

60. Jauniaux, J. – and Grenson, M. (1990). GAP1, the general amino acid permease gene of Saccharomyces cerevisiae Nucleotide sequence, protein similarity with the other bakers yeast amino acid permeases, and nitrogen catabolite repression. Eur J Biochem 190. 10.1111/j.1432-1033.1990.tb15542.x.

61. Schreve, J.L., Sin, J.K., and Garrett, J.M. (1998). The Saccharomyces cerevisiae YCC5 (YCL025c) gene encodes an amino acid permease, Agp1, which transports asparagine and glutamine. J Bacteriol 180. 10.1128/jb.180.9.2556-2559.1998.

62. Bianchi, F., van’t Klooster, J.S., Ruiz, S.J., and Poolman, B. (2019). Regulation of Amino Acid Transport in Saccharomyces cerevisiae. Microbiology and Molecular Biology Reviews 83. 10.1128/mmbr.00024-19.

63. Natarajan, K., Meyer, M.R., Jackson, B.M., Slade, D., Roberts, C., Hinnebusch, A.G., and Marton, M.J. (2001). Transcriptional Profiling Shows that Gcn4p Is a Master Regulator of Gene Expression during Amino Acid Starvation in Yeast. Mol Cell Biol 21. 10.1128/mcb.21.13.4347-4368.2001.

64. Yang, H., Tong, J., Lee, C.W., Ha, S., Eom, S.H., and Im, Y.J. (2015). Structural mechanism of ergosterol regulation by fungal sterol transcription factor Upc2. Nature Communications 2015 6:1 *6*, 1–13. 10.1038/ncomms7129.

65. White, W.H., Gunyuzlu, P.L., and Toyn, J.H. (2001). Saccharomyces cerevisiae Is Capable of de Novo Pantothenic Acid Biosynthesis Involving a Novel Pathway of β-Alanine Production from Spermine. Journal of Biological Chemistry 276. 10.1074/jbc.M009804200.

66. Boss, P.K., Pearce, A.D., Zhao, Y., Nicholson, E.L., Dennis, E.G., and Jeffery, D.W. (2015). Potential grape-derived contributions to volatile ester concentrations in wine. Molecules 20. 10.3390/molecules20057845.

67. Hanna, D. A., Harvey, R. M., Martinez-Guzman, O., Yuan, X., Chandrasekharan, B., Raju, G., Outten, F. W., Hamz, I., & Reddi, A. R. (2016). Heme dynamics and trafficking factors revealed by genetically encoded fluorescent heme sensors. Proceedings of the National Academy of Sciences of the United States of America, 113(27), 7539–7544. 10.1073/PNAS.1523802113

68. Breker, M., Gymrek, M., and Schuldiner, M. (2013). A novel single-cell screening platform reveals proteome plasticity during yeast stress responses. Journal of Cell Biology 200. 10.1083/jcb.201301120.

69. Schulte, U., den Brave, F., Haupt, A., Gupta, A., Song, J., Müller, C.S., Engelke, J., Mishra, S., Mårtensson, C., Ellenrieder, L., et al. (2023). Mitochondrial complexome reveals quality-control pathways of protein import. Nature 614. 10.1038/s41586-022-05641-w.

70. Wilson, J.D., and Barlowe, C. (2010). Yet1p and Yet3p, the yeast homologs of BAP29 and BAP31, interact with the endoplasmic reticulum translocation apparatus and are required for inositol prototrophy. Journal of Biological Chemistry 285, 18252–18261. 10.1074/jbc.M109.080382.

71. Altmann, K., and Westermann, B. (2005). Role of essential genes in mitochondrial morphogenesis in Saccharomyces cerevisiae. Mol Biol Cell 16. 10.1091/mbc.E05-07-0678.

72. Del Dedo, J.E., Fernández-Golbano, I., Pastor, L., Meler, P., Ferrer-Orta, C., Rebollo, E., and Geli, M.I. (2021). Coupled sterol synthesis and transport machineries at er–endocytic contact sites. Journal of Cell Biology 220. 10.1083/jcb.202010016.

73. Runz, H., Miura, K., Weiss, M., and Pepperkok, R. (2006). Sterols regulate ER-export dynamics of secretory cargo protein ts-O45-G. EMBO Journal 25. 10.1038/sj.emboj.7601205.

74. Lee, M., Moon, Y., Lee, S., Lee, C., and Jun, Y. (2019). Ergosterol interacts with Sey1p to promote atlastin-mediated endoplasmic reticulum membrane fusion in Saccharomyces cerevisiae. FASEB Journal 33. 10.1096/fj.201800779RR.

75. Lewis, P.M., Dunn, M.P., McMahon, J.A., Logan, M., Martin, J.F., St-Jacques, B., and McMahon, A.P. (2001). Cholesterol modification of sonic hedgehog is required for long-range signaling activity and effective modulation of signaling by Ptc1. Cell 105. 10.1016/S0092-8674(01)00369-5.

76. King, C., Sengupta, P., Seo, A.Y., and Lippincott-Schwartz, J. (2020). ER membranes exhibit phase behavior at sites of organelle contact. Proc Natl Acad Sci U S A 117. 10.1073/pnas.1910854117.

77. Kemmeren, P., Sameith, K., Van De Pasch, L.A.L., Benschop, J.J., Lenstra, T.L., Margaritis, T., O’Duibhir, E., Apweiler, E., Van Wageningen, S., Ko, C.W., et al. (2014). Large-scale genetic perturbations reveal regulatory networks and an abundance of gene-specific repressors. Cell 157. 10.1016/j.cell.2014.02.054.

78. Del Dedo, J.E., Fernández-Golbano, I., Pastor, L., Meler, P., Ferrer-Orta, C., Rebollo, E., and Geli, M.I. (2021). Coupled sterol synthesis and transport machineries at er–endocytic contact sites. Journal of Cell Biology 220. 10.1083/jcb.202010016.

79. Lee, M., Moon, Y., Lee, S., Lee, C., and Jun, Y. (2019). Ergosterol interacts with Sey1p to promote atlastin-mediated endoplasmic reticulum membrane fusion in Saccharomyces cerevisiae. FASEB Journal 33. 10.1096/fj.201800779RR.

80. Voeltz, G.K., Prinz, W.A., Shibata, Y., Rist, J.M., and Rapoport, T.A. (2006). A class of membrane proteins shaping the tubular endoplasmic reticulum. Cell 124. 10.1016/j.cell.2005.11.047.

81. Hu, J., Shibata, Y., Zhu, P.P., Voss, C., Rismanchi, N., Prinz, W.A., Rapoport, T.A., and Blackstone, C. (2009). A Class of Dynamin-like GTPases Involved in the Generation of the Tubular ER Network. Cell 138. 10.1016/j.cell.2009.05.025.

82. Cohen, N., Breker, M., Bakunts, A., Pesek, K., Chas, A., Argemí, J., Orsi, A., Gal, L., Chuartzman, S., Wigelman, Y., et al. (2017). Iron affects Ire1 clustering propensity and the amplitude of endoplasmic reticulum stress signaling. J Cell Sci 130, 3222–3233. 10.1242/JCS.201715.

83. Travers, K.J., Patil, C.K., Wodicka, L., Lockhart, D.J., Weissman, J.S., and Walter, P. (2000). Functional and genomic analyses reveal an essential coordination between the unfolded protein response and ER-associated degradation. Cell 101, 249–258. 10.1016/S0092-8674(00)80835-1.

84. Gasch, A.P., Spellman, P.T., Kao, C.M., Carmel-Harel, O., Eisen, M.B., Storz, G., Botstein, D., and Brown, P.O. (2000). Genomic expression programs in the response of yeast cells to environmental changes. Mol Biol Cell 11. 10.1091/mbc.11.12.4241.

85. Hulce, J.J., Cognetta, A.B., Niphakis, M.J., Tully, S.E., and Cravatt, B.F. (2013). Proteome-wide mapping of cholesterol-interacting proteins in mammalian cells. Nat Methods 10. 10.1038/nmeth.2368.

86. Jumper, J., Evans, R., Pritzel, A., Green, T., Figurnov, M., Ronneberger, O., Tunyasuvunakool, K., Bates, R., Žídek, A., Potapenko, A., et al. (2021). Highly accurate protein structure prediction with AlphaFold. Nature 596. 10.1038/s41586-021-03819-2.

87. Loewen, C.J.R., and Levine, T.P. (2005). A highly conserved binding site in vesicle-associated membrane protein-associated protein (VAP) for the FFAT motif of lipid-binding proteins. Journal of Biological Chemistry 280. 10.1074/jbc.M500147200.

88. Giaever, G., Chu, A.M., Ni, L., Connelly, C., Riles, L., Véronneau, S., Dow, S., Lucau-Danila, A., Anderson, K., André, B., et al. (2002). Functional profiling of the Saccharomyces cerevisiae genome. Nature 418. 10.1038/nature00935.

89. Breslow, D.K., Cameron, D.M., Collins, S.R., Schuldiner, M., Stewart-Ornstein, J., Newman, H.W., Braun, S., Madhani, H.D., Krogan, N.J., and Weissman, J.S. (2008). A comprehensive strategy enabling high-resolution functional analysis of the yeast genome. Nat Methods 5. 10.1038/nmeth.1234.

90. Gaspar, M.L., Hofbauer, H.F., Kohlwein, S.D., and Henry, S.A. (2011). Coordination of storage lipid synthesis and membrane biogenesis: Evidence for cross-talk between triacylglycerol metabolism and phosphatidylinositol synthesis. Journal of Biological Chemistry 286. 10.1074/jbc.M110.172296.

91. Cho, N.H., Cheveralls, K.C., Brunner, A.D., Kim, K., Michaelis, A.C., Raghavan, P., Kobayashi, H., Savy, L., Li, J.Y., Canaj, H., et al. (2022). OpenCell: Endogenous tagging for the cartography of human cellular organization. Science (1979) *375*. 10.1126/science.abi6983.

92. Iwasawa, R., Mahul-Mellier, A.L., Datler, C., Pazarentzos, E., and Grimm, S. (2011). Fis1 and Bap31 bridge the mitochondria-ER interface to establish a platform for apoptosis induction. EMBO Journal 30. 10.1038/emboj.2010.346.

93. Cho, N.H., Cheveralls, K.C., Brunner, A.D., Kim, K., Michaelis, A.C., Raghavan, P., Kobayashi, H., Savy, L., Li, J.Y., Canaj, H., et al. (2022). OpenCell: Endogenous tagging for the cartography of human cellular organization. Science (1979) *375*. 10.1126/science.abi6983.

94. Wei, X., Li, L., Zhao, J., Huo, Y., Hu, X., Lu, J., Pi, J., Zhang, W., Xu, L., Yao, Y., et al. (2023). BAP31 depletion inhibited adipogenesis, repressed lipolysis and promoted lipid droplets abnormal growth via attenuating Perilipin1 proteasomal degradation. Int J Biol Sci 19. 10.7150/ijbs.82178.

95. Luo, W., Adamska, J.Z., Li, C., Verma, R., Liu, Q., Hagan, T., Wimmers, F., Gupta, S., Feng, Y., Jiang, W., et al. (2023). SREBP signaling is essential for effective B cell responses. Nat Immunol 24. 10.1038/s41590-022-01376-y.

96. Geisberger, R., Lamers, M., & Achatz, G. (2006). The riddle of the dual expression of IgM and IgD. In Immunology (Vol. 118, Issue 4). 10.1111/j.1365-2567.2006.02386.x

97. Paquet, M.-E., Cohen-Doyle, M., Shore, G.C., and Williams, D.B. (2004). Bap29/31 Influences the Intracellular Traffic of MHC Class I Molecules. The Journal of Immunology 172. 10.4049/jimmunol.172.12.7548.

98. Niu, K., Xu, J., Cao, Y., Hou, Y., Shan, M., Wang, Y., Xu, Y., Sun, M., and Wang, B. (2017). BAP31 is involved in T cell activation through TCR signal pathways. Sci Rep 7. 10.1038/srep44809.

99. Drake, D.R., and Braciale, T.J. (2001). Cutting Edge: Lipid Raft Integrity Affects the Efficiency of MHC Class I Tetramer Binding and Cell Surface TCR Arrangement on CD8+ T Cells. The Journal of Immunology 166. 10.4049/jimmunol.166.12.7009.

100. Zhao, B., Sun, L., Yuan, Q., Hao, Z., An, F., Zhang, W., Zhu, X., and Wang, B. (2023). BAP31 Knockout in Macrophages Affects CD4+T Cell Activation through Upregulation of MHC Class II Molecule. Int J Mol Sci 24. 10.3390/ijms241713476.

101. Xu, K., Han, B., Bai, Y., Ma, X.Y., Ji, Z.N., Xiong, Y., Miao, S.K., Zhang, Y.Y., and Zhou, L.M. (2019). MiR-451a suppressing BAP31 can inhibit proliferation and increase apoptosis through inducing ER stress in colorectal cancer. Cell Death & Disease 2019 10:3 *10*, 1–16. 10.1038/s41419-019-1403-x.

102. Wang, A., Zhang, Y., and Cao, P. (2019). Inhibition of BAP31 expression inhibits cervical cancer progression by suppressing metastasis and inducing intrinsic and extrinsic apoptosis. Biochem Biophys Res Commun 508, 499–506. 10.1016/J.BBRC.2018.11.017.

103. Chen, J., Guo, H., Jiang, H., Namusamba, M., Wang, C., Lan, T., Wang, T., and Wang, B. (2019). A BAP31 intrabody induces gastric cancer cell death by inhibiting p27kip1 proteasome degradation. Int J Cancer 144, 2051–2062. 10.1002/IJC.31930.

104. Yu, S., Wang, F., Fan, L., Wei, Y., Li, H., Sun, Y., Yang, A., Jin, B., Song, C., and Yang, K. (2015). BAP31, a promising target for the immunotherapy of malignant melanomas. J Exp Clin Cancer Res 34. 10.1186/S13046-015-0153-6.

105. Liu, X., Jiao, K., Jia, C. cong, Li, G. xun, Yuan, Q., Xu, J. kai, Hou, Y., and Wang, B. (2019). BAP31 regulates IRAK1-dependent neuroinflammation in microglia. J Neuroinflammation 16. 10.1186/S12974-019-1661-7.

106. Wang, T., Chen, J., Hou, Y., Yu, Y., and Wang, B. (2019). BAP31 deficiency contributes to the formation of amyloid-β plaques in Alzheimer’s disease by reducing the stability of RTN3. FASEB J 33, 4936–4946. 10.1096/FJ.201801702R.

107. Regan, J.A., and Laimins, L.A. (2008). Bap31 is a novel target of the human papillomavirus E5 protein. J Virol 82, 10042–10051. 10.1128/JVI.01240-08.

108. Geiger, R., Andritschke, D., Friebe, S., Herzog, F., Luisoni, S., Heger, T., and Helenius, A. (2011). BAP31 and BiP are essential for dislocation of SV40 from the endoplasmic reticulum to the cytosol. Nat Cell Biol 13, 1305–1314. 10.1038/NCB2339.

109. Brachmann, C.B., Davies, A., Cost, G.J., Caputo, E., Li, J., Hieter, P., and Boeke, J.D. (1998). Designer deletion strains derived from Saccharomyces cerevisiae S288C: A useful set of strains and plasmids for PCR-mediated gene disruption and other applications. Yeast 14. 10.1002/(SICI)1097-0061(19980130)14:2<115::AID-YEA204>3.0.CO;2-2.

110. Gietz, R.D., and Woods, R.A. (2006). Yeast transformation by the LiAc/SS Carrier DNA/PEG method. Methods Mol Biol 313. 10.1385/1-59259-958-3:107.

111. Janke, C., Magiera, M.M., Rathfelder, N., Taxis, C., Reber, S., Maekawa, H., Moreno-Borchart, A., Doenges, G., Schwob, E., Schiebel, E., et al. (2004). A versatile toolbox for PCR-based tagging of yeast genes: New fluorescent proteins, more markers and promoter substitution cassettes. Yeast 21. 10.1002/yea.1142.

112. Longtine, M.S., McKenzie, A., Demarini, D.J., Shah, N.G., Wach, A., Brachat, A., Philippsen, P., and Pringle, J.R. (1998). Additional modules for versatile and economical PCR-based gene deletion and modification in Saccharomyces cerevisiae. Yeast 14. 10.1002/(SICI)1097-0061(199807)14:10<953::AID-YEA293>3.0.CO;2-U.

113. Yofe, I., and Schuldiner, M. (2014). Primers-4-Yeast: A comprehensive web tool for planning primers for Saccharomyces cerevisiae. Yeast 31. 10.1002/yea.2998.

114. Schindelin, J., Arganda-Carreras, I., Frise, E., Kaynig, V., Longair, M., Pietzsch, T., Preibisch, S., Rueden, C., Saalfeld, S., Schmid, B., Tinevez, J. Y., White, D. J., Hartenstein, V., Eliceiri, K., Tomancak, P., & Cardona, A. (2012). Fiji: An open-source platform for biological-image analysis. In Nature Methods (Vol. 9, Issue 7). 10.1038/nmeth.2019

115. Mastronarde, D.N. (2005). Automated electron microscope tomography using robust prediction of specimen movements. J Struct Biol 152. 10.1016/j.jsb.2005.07.007.

116. Kremer, J.R., Mastronarde, D.N., and McIntosh, J.R. (1996). Computer visualization of three-dimensional image data using IMOD. J Struct Biol 116. 10.1006/jsbi.1996.0013.

117. Hagen, W.J.H., Wan, W., and Briggs, J.A.G. (2017). Implementation of a cryo-electron tomography tilt-scheme optimized for high resolution subtomogram averaging. J Struct Biol 197. 10.1016/j.jsb.2016.06.007.

118. Mastronarde, D.N., and Held, S.R. (2017). Automated tilt series alignment and tomographic reconstruction in IMOD. J Struct Biol 197. 10.1016/j.jsb.2016.07.011.

119. Liu, Y.T., Zhang, H., Wang, H., Tao, C.L., Bi, G.Q., and Zhou, Z.H. (2022). Isotropic reconstruction for electron tomography with deep learning. Nat Commun 13. 10.1038/s41467-022-33957-8.

120. Lamm, L., Zufferey, S., Righetto, R.D., Wietrzynski, W., Yamauchi, K.A., Burt, A., Liu, Y., Zhang, H., Martinez-Sanchez, A., Ziegler, S., et al. (2024). MemBrain v2: an end-to-end tool for the analysis of membranes in cryo-electron tomography. bioRxiv.

121. Pettersen, E.F., Goddard, T.D., Huang, C.C., Couch, G.S., Greenblatt, D.M., Meng, E.C., and Ferrin, T.E. (2004). UCSF Chimera – A visualization system for exploratory research and analysis. J Comput Chem 25. 10.1002/jcc.20084.

122. Tong, A.H.Y., and Boone, C. (2006). Synthetic genetic array analysis in Saccharomyces cerevisiae. Methods Mol Biol 313. 10.1385/1-59259-958-3:171.

123. Breslow, D.K., Cameron, D.M., Collins, S.R., Schuldiner, M., Stewart-Ornstein, J., Newman, H.W., Braun, S., Madhani, H.D., Krogan, N.J., and Weissman, J.S. (2008). A comprehensive strategy enabling high-resolution functional analysis of the yeast genome. Nat Methods 5. 10.1038/nmeth.1234.

124. Perez-Riverol, Y., Bai, J., Bandla, C., García-Seisdedos, D., Hewapathirana, S., Kamatchinathan, S., Kundu, D.J., Prakash, A., Frericks-Zipper, A., Eisenacher, M., et al. (2022). The PRIDE database resources in 2022: A hub for mass spectrometry-based proteomics evidences. Nucleic Acids Res 50. 10.1093/nar/gkab1038.

125. Elinger, D., Gabashvili, A., and Levin, Y. (2019). Suspension Trapping (S-Trap) Is Compatible with Typical Protein Extraction Buffers and Detergents for Bottom-Up Proteomics. J Proteome Res 18. 10.1021/acs.jproteome.8b00891.

126. Cox, J., Hein, M.Y., Luber, C.A., Paron, I., Nagaraj, N., and Mann, M. (2014). Accurate proteome-wide label-free quantification by delayed normalization and maximal peptide ratio extraction, termed MaxLFQ. Molecular and Cellular Proteomics 13. 10.1074/mcp.M113.031591.

127. Tyanova, S., Temu, T., Sinitcyn, P., Carlson, A., Hein, M. Y., Geiger, T., Mann, M., & Cox, J. (2016). The Perseus computational platform for comprehensive analysis of (prote)omics data. In Nature Methods (Vol. 13, Issue 9). 10.1038/nmeth.3901

128. Malitsky, S., Ziv, C., Rosenwasser, S., Zheng, S., Schatz, D., Porat, Z., Ben-Dor, S., Aharoni, A., and Vardi, A. (2016). Viral infection of the marine alga Emiliania huxleyi triggers lipidome remodeling and induces the production of highly saturated triacylglycerol. New Phytologist 210. 10.1111/nph.13852.

129. Zheng, L., Cardaci, S., Jerby, L., Mackenzie, E.D., Sciacovelli, M., Johnson, T.I., Gaude, E., King, A., Leach, J.D.G., Edrada-Ebel, R., et al. (2015). Fumarate induces redox-dependent senescence by modifying glutathione metabolism. Nat Commun 6. 10.1038/ncomms7001.

130. Dong, Y., Steffenson, B.J., and Mirocha, C.J. (2006). Analysis of ergosterol in single kernel and ground grain by gas chromatography-mass spectrometry. J Agric Food Chem 54. 10.1021/jf060149f.

131. Zhang, Y., Karmon, O., Das, K., Wiener, R., Lehming, N., and Pines, O. (2022). Ubiquitination Occurs in the Mitochondrial Matrix by Eclipsed Targeted Components of the Ubiquitination Machinery. Cells 11. 10.3390/cells11244109.

132. Martinez-Guzman, O., Willoughby, M.M., Saini, A., Dietz, J. V., Bohovych, I., Medlock, A.E., Khalimonchuk, O., and Reddi, A.R. (2020). Mitochondrial-nuclear heme trafficking in budding yeast is regulated by GTPases that control mitochondrial dynamics and ER contact sites. J Cell Sci 133. 10.1242/jcs.237917.

133. Hanna, D.A., Moore, C.M., Liu, L., Yuan, X., Dominic, I.M., Fleischhacker, A.S., Hamza, I., Ragsdale, S.W., and Reddi, A.R. (2022). Heme oxygenase-2 (HO-2) binds and buffers labile ferric heme in human embryonic kidney cells. Journal of Biological Chemistry 298. 10.1016/j.jbc.2021.101549.

