## Supplementary figures and images for "The molecular mechanism of on-demand sterol biosynthesis at organelle contact sites"

### Figure S1

Zung et al, Figure S1

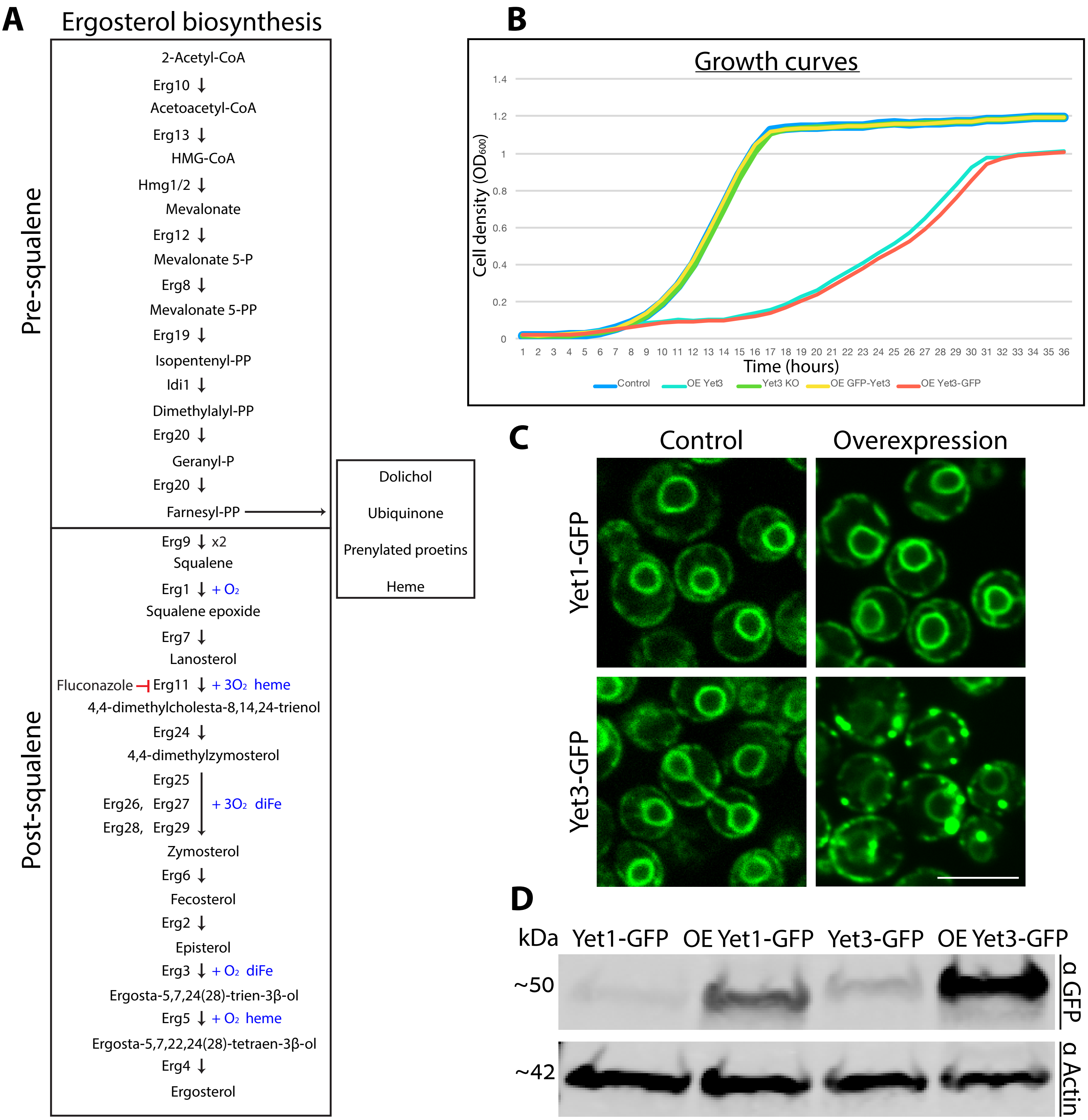

### Figure S2

Zung et al, Figure S2

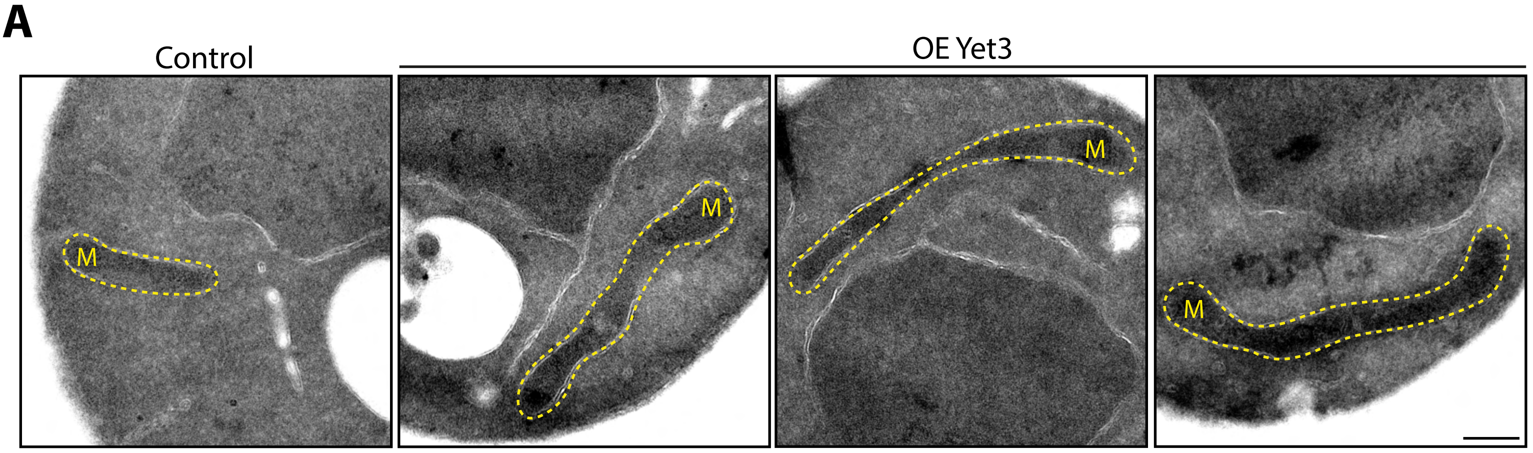


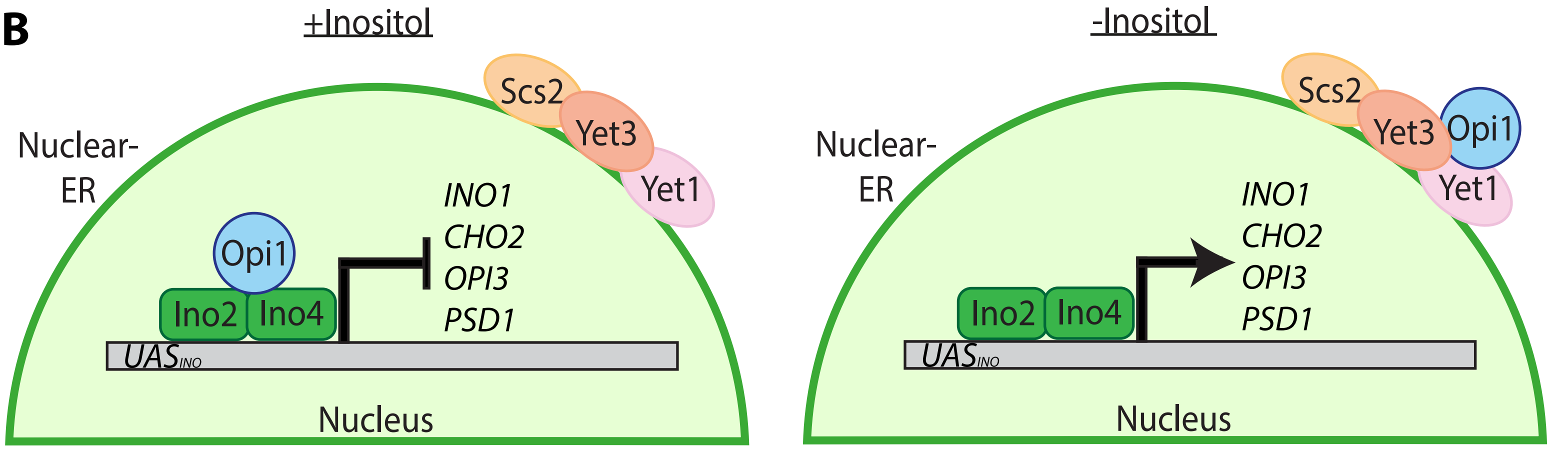

### Figure S3

Zung et al, Figure S3

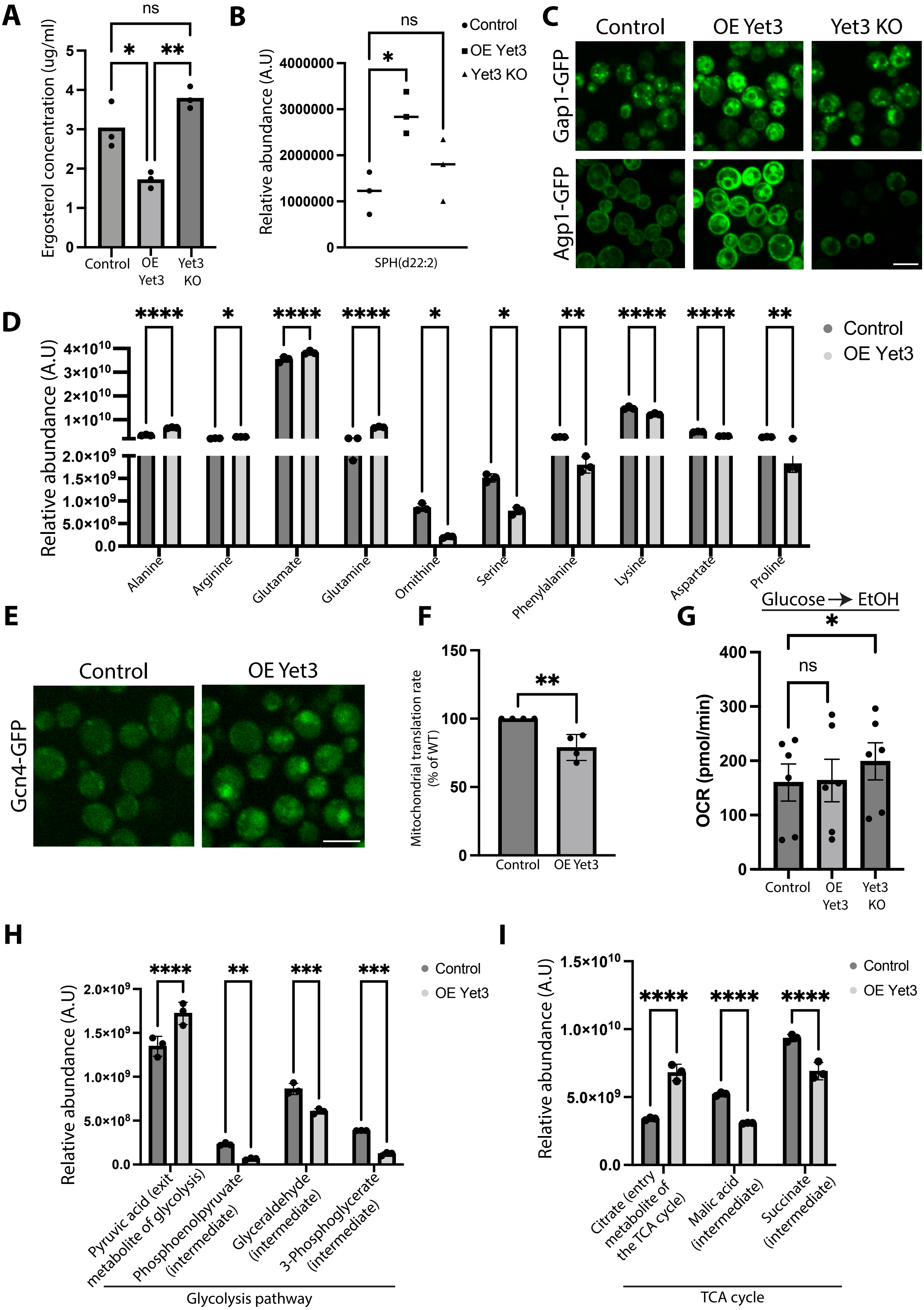

### Figure S4A

Zung et al, Figure S4- A

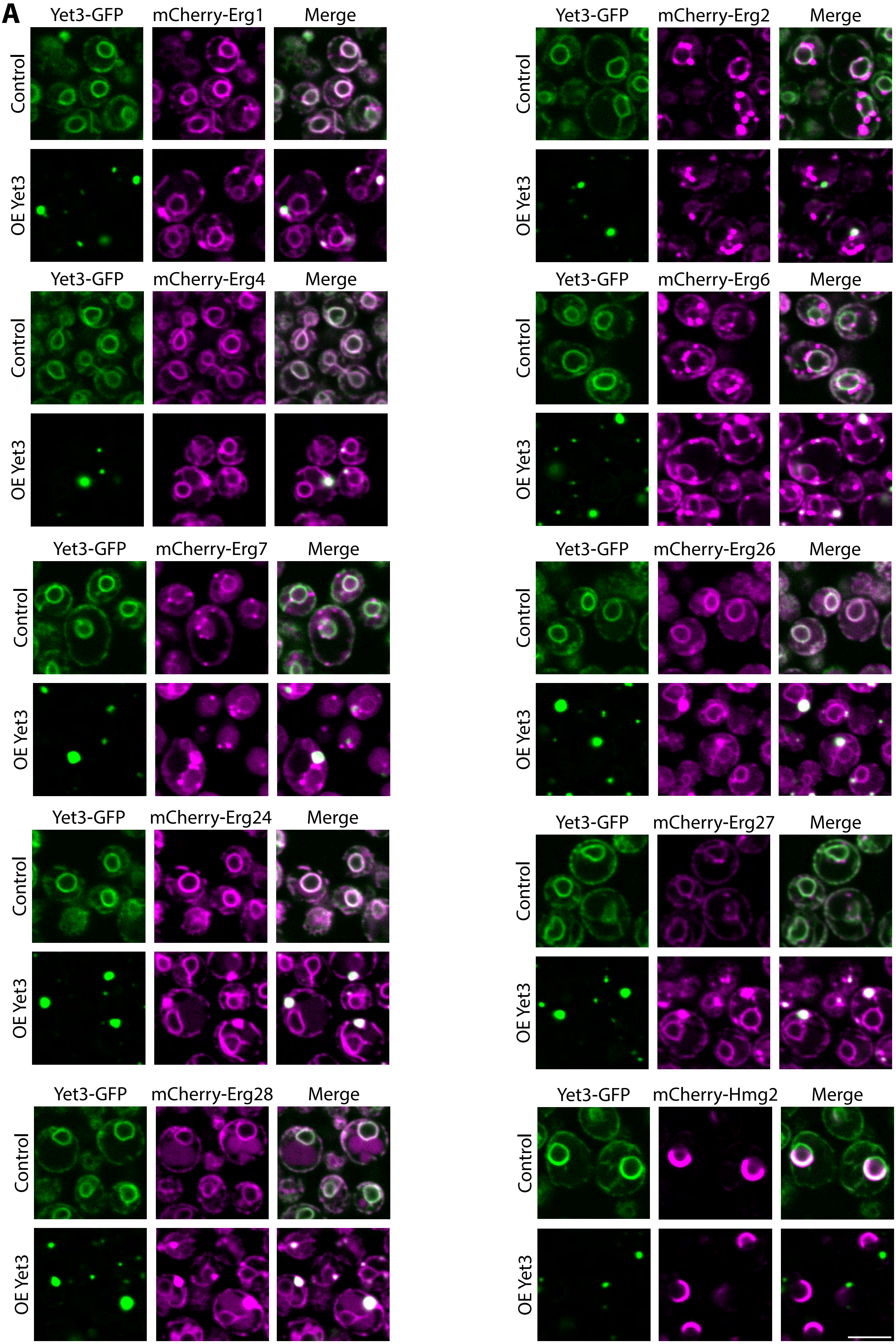

### Figure S4B-F

Zung et al, Figure S4 B-F

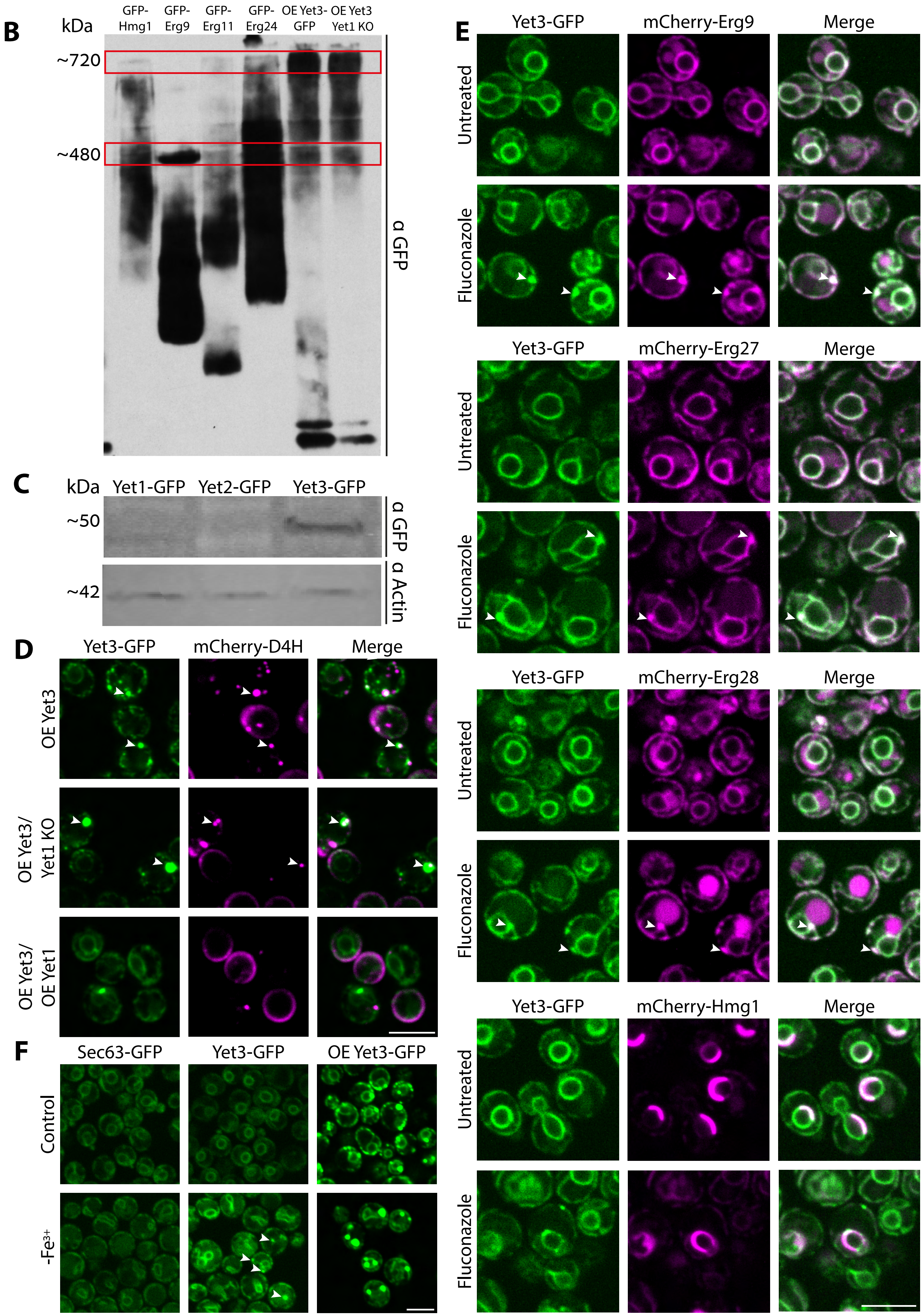

### Figure S5

Zung et al, Figure S5

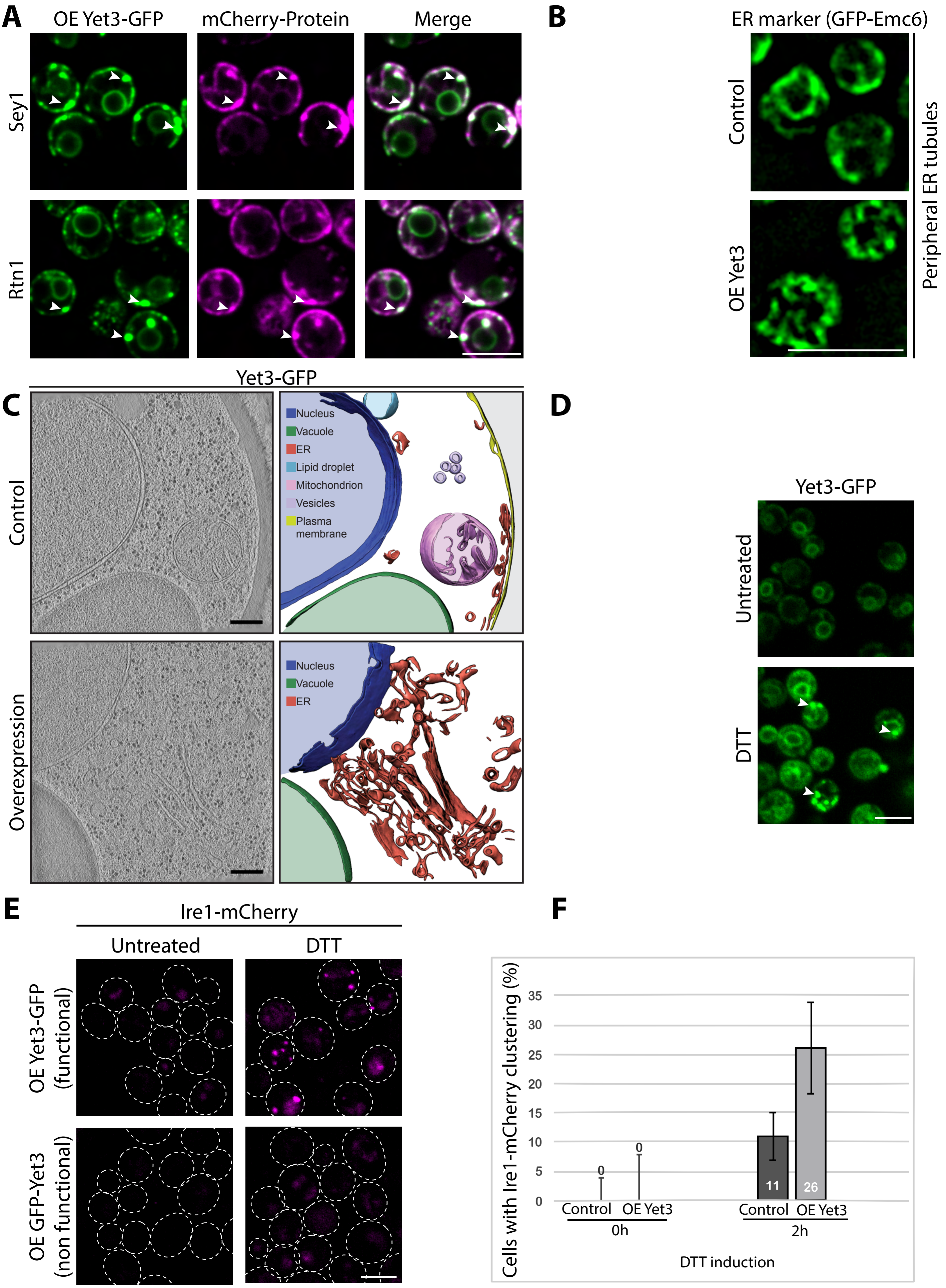

### Figure S6

Zung et al, Figure S6

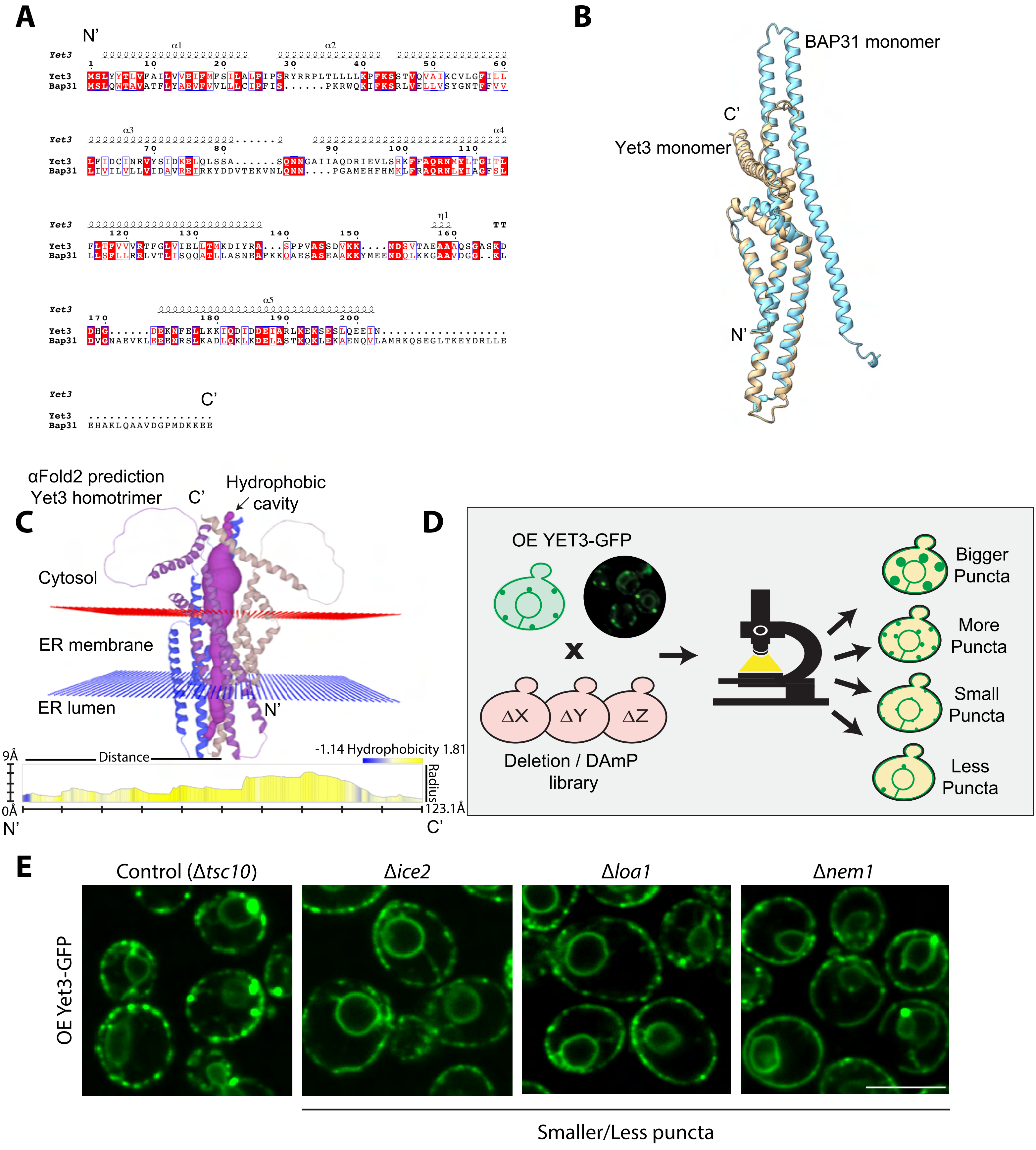
